# Parthenolide Covalently Targets and Inhibits Focal Adhesion Kinase in Breast Cancer Cells

**DOI:** 10.1101/550806

**Authors:** Charles A. Berdan, Raymond Ho, Haley S. Lehtola, Milton To, Xirui Hu, Tucker R. Huffman, Yana Petri, Chad R. Altobelli, Sasha G. Demeulenaere, James A. Olzmann, Thomas J. Maimone, Daniel K. Nomura

## Abstract

Parthenolide, a natural product from the feverfew plant and member of the large family of sesquiterpene lactones, exerts multiple biological and therapeutic activities including anti-inflammatory and anti-cancer effects. Herein, we further study parthenolide mechanism of action using activity-based protein profiling (ABPP)-based chemoproteomic platforms to map additional covalent targets engaged by parthenolide in human breast cancer cells. We find that parthenolide, as well as other related exocyclic methylene lactone-containing sesquiterpenes, covalently modify cysteine 427 (C427) of focal adhesion kinase 1 (FAK1) leading to impairment of FAK1-dependent signaling pathways and breast cancer cell proliferation, survival, and motility. These studies reveal a novel functional target exploited by members of a large family of anticancer natural products.

## Main text

Parthenolide, a natural product found in the feverfew plant (*Tanacetum parthenium*), possesses myriad therapeutic activities, including anti-inflammatory and anti-cancer effects. Through covalent bond formation between its reactive α-methylene-γ-butyrolactone moiety and various protein targets, multiple cellular signaling pathways are impacted (Ghantous et al., 2013; Kwok et al., 2001; Liu et al., 2018; Mathema et al., 2012; Shin et al., 2017). Moreover, this natural product belongs to the broader family of sesquiterpene lactones (estimated at > 5000 members), many members of which are also cytotoxic and have been hypothesized or shown to act through covalent mechanisms (Coricello et al., 2018; Quintana and Estévez, 2019). Parthenolide impairs cancer pathogenicity or confers chemotherapy or radiation sensitivity across a wide range of cancer types, including leukemia, colorectal, glioblastoma, cervical, liver, prostate, lung, pancreatic, skin, and breast cancers (Anderson and Bejcek, 2008; Carlisi et al., 2016; Diamanti et al., 2013; Jeyamohan et al., 2016; Kim et al., 2012, 2017; Lesiak et al., 2010; Lin et al., 2017; Liu et al., 2017; Morel et al., 2017; Ralstin et al., 2006; Sun et al., 2007; Sweeney et al., 2005). Despite possessing multi-target activity and exhibiting cytotoxicity across a wide range of human cancers, parthenolide is remarkably well-tolerated in humans (Curry et al., 2004).

Using a biotinylated parthenolide analog, previous studies by the lab of Crews established that one of the primary targets that drives the anti-inflammatory and anti-cancer activity of parthenolide is IKK-β wherein cysteine 179 (C179) is modified thus impairing IKK-β and NFκB signaling (Kwok et al., 2001). Additional studies have revealed other direct targets of parthenolide that may help to explain the therapeutic properties of this natural product, including targeting of specific cysteines within heat shock protein Hsp72 and STAT3 downstream signaling targets such as Janus kinases JAK2 (Liu et al., 2018; Shin et al., 2017). Moreover, this natural product has also been shown to affect additional cell signaling pathways including induction of oxidative stress and apoptosis, focal adhesion kinase 1 (FAK1) signaling, HIF-1α signaling, epithelial-to-mesenchymal transition, Wnt/β-catenin signaling, MAPK signaling, and mitochondrial function (Carlisi et al., 2011, 2016; Jafari et al., 2018; Kim et al., 2017; Kwok et al., 2001; Lin et al., 2017; Zhang et al., 2017). Based on the broader scope of influence on these biological pathways and systems, parthenolide likely still possesses additional targets that are not yet fully elucidated. In previous works investigating the direct targets of parthenolide, multiple studies have revealed unique ligandable and functional cysteines within their respective proteins that could be targeted to influence cellular signaling and pathogenicity. Recent studies have shown that activity-based protein profiling (ABPP)-based chemoproteomic platforms can be utilized to uncover unique and functional druggable hotspots and modalities that can be accessed by covalently-acting small-molecules and natural products that may not be obvious using standard drug discovery paradigms (Backus et al., 2016; Bateman et al., 2017; Grossman et al., 2017; Hacker et al., 2017; Spradlin et al., 2018; Ward et al., 2018). ABPP uses reactivity-based chemical probes to profile proteome-wide reactive, ligandable, and functional sites directly in complex proteomes. When used in a competitive manner, covalently-acting small-molecules can be competed against binding of reactivity-based probes to map the proteome-wide targets of these compounds (Backus et al., 2016; Bateman et al., 2017; Grossman et al., 2017; Hacker et al., 2017; Roberts et al., 2017a; Wang et al., 2014). Importantly, this technology allows for the interrogation of natural products in their unmodified form.

In this study, we used ABPP chemoproteomic platforms to map additional targets of parthenolide in breast cancer cells uncovering additional druggable hotspots which may contribute to the cell signaling and anti-cancer effects of parthenolide (Fig. 1A). Parthenolide impaired cell survival and proliferation, induced apoptosis, thwarted early cell motility, and significantly attenuated *in vivo* tumor xenograft growth in estrogen receptor, progesterone receptor, and HER2 receptor-negative breast cancer (triple-negative breast cancer, TNBC) cells, 231MFP or HCC38 (Fig. 1B-1F). We note that we are observing anti-tumorigenic effects at a relatively low dose of 30 mg/kg, despite observing cell viability impairments at 50 μM high concentrations. This may be because of the covalent nature of parthenolide and accumulating target engagement over time. Since parthenolide irreversibly binds to their targets, the targets will stay bound to parthenolide until the protein turns over. TNBCs are amongst the most aggressive of breast cancer subtypes; they show the worst prognoses and currently have few targeted therapies. Our data suggested that parthenolide may be effective at attenuating TNBC pathogenicity.

**Figure 1.**
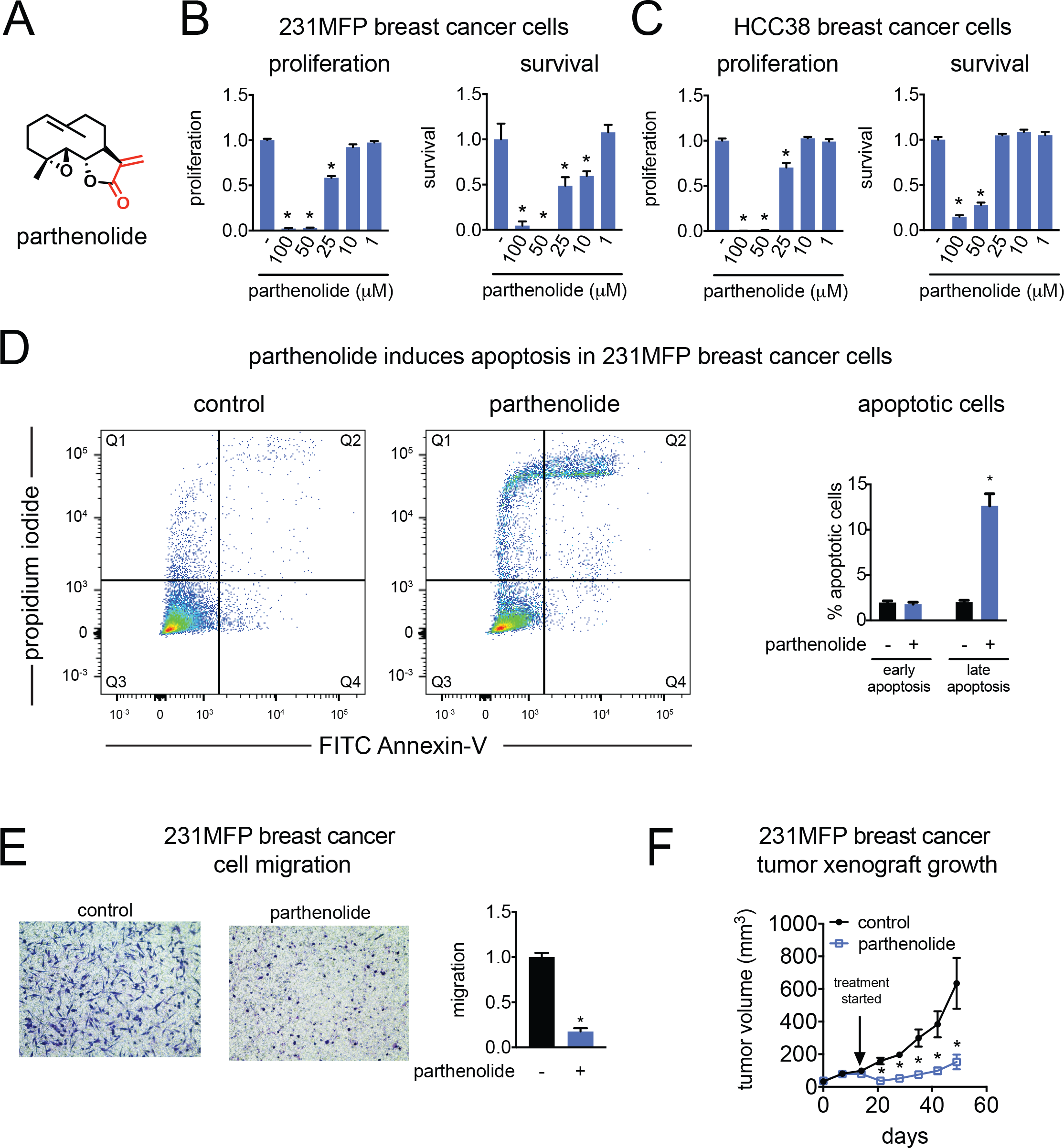
Parthenolide impairs TNBC pathogenicity. **(A)** Structure of parthenolide (red denotes cysteine-reactive enone). **(B-C)** 231MFP **(B)** and HCC38 **(C)** breast cancer cell 48 h survival and proliferation from cells treated with DMSO control or parthenolide assessed by Hoechst stain. **(D)** Percent early-stage and late-stage apoptotic cells assessed by flow cytometry after treating cells with DMSO control or parthenolide (50 μM) for 2 h. Shown on the left panels are representative FACS data. On the right bar graph are noted early apoptotic cells defined as FITC+/PI- and late apoptotic cells defined as FITC+/PI+ cells. **(E)** Migration of 231MFP cells treated with DMSO control or parthenolide (50 μM) for 6 h. Representative image of migrated cells are shown. **(F)** 231MFP tumor xenograft growth in immune-deficient SCID mice treated with vehicle (18:1:1 saline:ethanol:PEG40) or parthenolide (30 mg/kg ip) daily once per day with treatment initiated after tumor establishment 14 days after tumor implantation. Images shown in **(E)** are representative of n=3. Data shown in **(B-F)** are average ± sem, n=3-6/group. Significance is expressed as *p<0.05 compared to vehicle-treated controls.

We next used ABPP methods to identify additional targets of parthenolide in breast cancer cells. To confirm that parthenolide was not completely non-specific, we first performed a competitive gel-based ABPP experiment in which we competed parthenolide against labeling of 231MFP breast cancer cell proteomes with a rhodamine-functionalized cysteine-reactive iodoacetamide (IA-rhodamine) probe. While this method is imprecise, we observed that parthenolide did not broadly inhibit global proteome-wide cysteine reactivity **(Fig. S1).** Using a more specific, previously-reported alkyne-functionalized parthenolide probe (parthenolide-alkyne) (Shin et al., 2017), we observed multiple labeled proteins in 231MFP proteomes, of which some, but not all, targets were competed by parthenolide **(Fig. S1)**. Collectively, these results indicated that parthenolide does possess multiple protein targets in 231MFP proteomes, but that this natural product is not completely promiscuous in its reactivity.

While the parthenolide-alkyne probe could be used to identify additional targets of this natural product, we sought to map the specific amino acids within these targets that were engaged by unfunctionalized parthenolide. Thus, we next used isotopic tandem orthogonal proteolysis-enabled ABPP (isoTOP-ABPP) to identify specific ligandable sites targeted by parthenolide in 231MFP breast cancer proteomes. We competed parthenolide binding against the broadly cysteine-reactive alkyne-functionalized iodoacetamide probe (iodoacetamide-alkyne, IA-alkyne) directly in 231MFP TNBC proteomes using previously established methods (Fig. 2A, **Table S1**) (Backus et al., 2016; Bateman et al., 2017; Grossman et al., 2017; Roberts et al., 2017a; Weerapana et al., 2010). This analysis revealed three highly engaged targets of parthenolide that showed isotopically light vehicle-treated to heavy parthenolide-treated probe-modified peptide ratios of greater than 10, indicating >90 % engagement of these sites—focal adhesion kinase 1 (FAK1) C427, paraoxonase 3 (PON3) C240, and DNA-protein kinase (DNA-PK or PRKDC) C729. FAK1 C427 was the top target showing the highest ratio, and thus we placed subsequent focus on investigating the role of FAK1-dependent effects of parthenolide in breast cancer cells (Fig. 2A; **Table S1**). While the role of PON3 in cancer cells is unclear, FAK1 and PRKDC are known to be important drivers of cancer cell signaling and DNA repair, respectively. Notably, FAK1 and DNA-PK inhibitors have been shown to impair cell proliferation and viability in cancer cells and are being pursued in the clinic (Helleday et al., 2008; Sulzmaier et al., 2014; Lv et al. 2018). Since C427 of FAK1 was the most highly engaged target in this study, we focused our attention on investigating the FAK1-dependent effects of parthenolide in TNBC cells.

**Figure 2.**
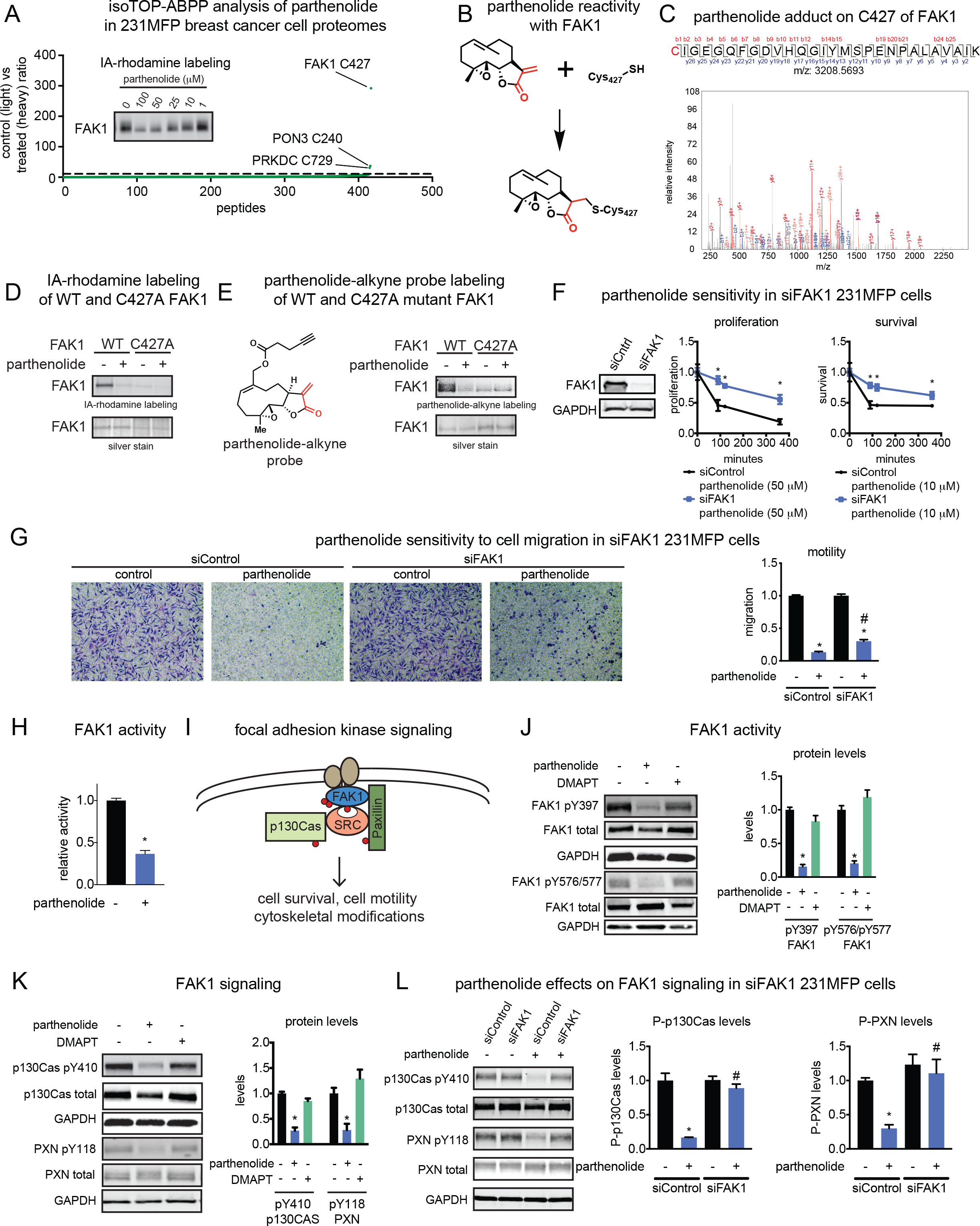
Parthenolide targets an allosteric cysteine in FAK1 to inhibit FAK1 activity and signaling. **(A)** isoTOP-ABPP analysis of parthenolide in 231MFP cell lysate. 231MFP proteomes were pre-treated with DMSO vehicle or parthenolide (50 μM) for 30 min prior to IA-alkyne (100 μM) labeling for 1 h. Shown are probe-modified peptides detected in at least two of out four biological replicates and the control (isotopically light) versus parthenolide-treated (isotopically heavy) ratios for each peptide. Those probe-modified peptides showing ratios >10 are peptides that were observed in three out of four biological replicates. Shown in the inset is gel-based ABPP analysis of parthenolide competition against IA-rhodamine labeling of pure human FAK1 kinase domain. **(B)** Proposed reactivity of parthenolide with C427 of FAK1. **(C)** Mass spectrometry data of covalent parthenolide adduct on C427 of human FAK1 protein. Pure human FAK1 kinase domain was labeled with parthenolide (100 μM) for 30 minutes and subsequently digested with trypsin for LC-MS/MS proteomic analysis. **(D)** Gel-based ABPP analysis of recombinant human wild-type and C427A mutant FAK1 kinase domain protein pre-incubated with parthenolide (50 μM) for 30 min prior to IA-rhodamine labeling (1 μM) for 1 h. Proteins were separated by SDS/PAGE and visualized by in-gel fluorescence. Also shown is silver-staining of the gel as a loading control. **(E)** Parthenolide-alkyne probe labeling of recombinant human wild-type and C427A mutant FAK1 kinase domain protein pre-incubated with parthenolide (50 μM) for 30 min prior to parthenolide-alkyne labeling (50 μM) for 1h. Rhodamine-azide was appended to probe-labeled protein by CuAAC and proteins were separated by SDS/PAGE and visualized by in-gel fluorescence. Also shown is silver-staining of the gel as a loading control. **(F)** FAK1 knockdown in 231MFP cells using siControl or siFAK1 oligonucleotides confirmed by Western blotting. Serum-containing cell proliferation and serum-free cell survival in siControl or siFAK1 231MFP cells treated with parthenolide (10 μM) for the reported time period. **(G)** Cell migration in siControl or siFAK1 231MFP cells treated with parthenolide (10 μM) for 4 h. **(H)** FAK1 activity with pure human FAK1 protein pre-treated with vehicle DMSO or parthenolide (100 μM) for 30 min before addition of peptide substrate and ATP. Activity was assessed by substrate peptide phosphorylation reading out ADP release using an ADP-Glo kinase assay. **(I)** FAK1 signaling pathways. **(J-K)** FAK1 signaling assessed by Western blotting in 231MFP cells treated with vehicle DMSO or parthenolide (50 μM) for 2 h. **(L)** FAK1 signaling assessed by Western blotting in siControl and siFAK1 cells treated with vehicle DMSO or parthenolide (50 μM) for 2 h. Gels shown in **(A, D-F, J-L)** are representative images from n=3 biological replicates/group. Data shown in **(F-H, J-L)** are average ± sem, n=3-6/group. Significance is expressed as *p<0.05 compared to vehicle-treated controls, # comparing parthenolide-treated siFAK1 cells compared to parthenolide-treated siControl cells. **Figure 2** is related to **Table S1** and **Figure S1**.

We validated the interaction of parthenolide with C427 of FAK1 using several complementary approaches. We first validated the interaction of parthenolide with FAK1 in which we showed parthenolide prevention of pure human FAK1 kinase domain cysteine-reactivity with a rhodamine-functionalized iodoacetmide probe (IA-rhodamine) by gel-based ABPP (Fig. 2A). Based on previous studies, we conjectured that parthenolide reacted covalently with C427 of FAK1 through a homo-Michael addition involving the α/β unsaturated lactone (Fig. 2B) (Kwok et al., 2001). Second, we demonstrated that parthenolide covalently reacts with C427 of FAK1 by identifying this parthenolide adduct on human FAK1 kinase domain by LC-MS/MS (Fig. 2C). We also demonstrated that IA-rhodamine labeling of pure human FAK1 was abrogated in the C427A mutant and that no additional inhibition of remaining IA-rhodamine labeling of FAK1 was observed with parthenolide treatment (Fig. 2D). Using a parthenolide-alkyne probe, we further showed that this probe labeled wild-type FAK1 protein, and that this labeling was prevented by parthenolide or in the C427A mutant FAK1 protein (Fig. 2E).

Previous studies have shown that FAK1 is amplified or overexpressed across a large fraction of breast tumors wherein FAK1 activity and expression is correlated with poor prognosis. FAK1 has been shown to be important in breast cancer cell survival, proliferation, and migration (Luo and Guan, 2010; Sulzmaier et al., 2014). To determine whether any of the observed parthenolide-mediated proliferative, survival, or migration impairments were dependent on FAK1, we assessed parthenolide effects on these phenotypes under FAK1 knockdown in 231MFP breast cancer cells (Fig. 2F-2G). FAK1 knockdown confers significant resistance to parthenolide-mediated impairments in cell proliferation, serum-free cell survival, and cell migration, particularly at early time-points, compared to control cells (Fig. 2F-2G), demonstrating that FAK1 contributes to the anti-cancer effects of parthenolide. Because parthenolide rapidly impairs cell proliferation and survival, we do note that the migration phenotypes shown here are likely confounded by reduced cell viability from parthenolide treatment. Interestingly, FAK1 knockdown by small-interfering RNA (siRNA) did not impair basal cell proliferation, survival, or migration. We postulate that this lack of effect may either be due to the multi-target polypharmacological nature of parthenolide, or potential adaptation to FAK1 knockdown during the inherently slower process of siRNA-mediated knockdown compared to acute inhibition of FAK1. We later show evidence for the latter hypothesis.

We next sought to determine whether parthenolide functionally inhibits FAK1 activity and signaling. Based on previously reported crystal structures of FAK1, C427 resided in a loop region proximal to the ATP site, indicating that covalent modification of this site may be inhibitory (Iwatani et al., 2013). Consistent with this premise, we showed that FAK1 activity was inhibited by parthenolide *in vitro* with pure human FAK1 kinase domain in a substrate activity assay (Fig. 2H). While this manuscript was under revision, an elegant study describing the first structure-guided design, synthesis, and characterization of a FAK1 inhibitor that also covalently targeted C427 of FAK1 and inhibited its function was reported (Yen-Pon et al., 2018). Importantly, this report gives further credence to our hypothesis of the functional relevance of this cysteine and its effects on cancer cell proliferation.

FAK1 is activated through membrane recruitment by growth factors, extracellular matrix, and integrin signaling followed by subsequent autophosphorylation at Y397. This produces an SH2-binding domain, which in-turn recruits Src and promotes semi-autophosphorylation of Y576/577 of FAK1. The fully active FAK1/Src complex can now recruit, phosphorylate, and activate numerous targets including p130Cas/Bcar1 and paxillin (PXN) to drive cell motility and cytoskeletal modifications (Frame et al., 2010; Sulzmaier et al., 2014) (Fig. 2I). We show that parthenolide, but not the analog dimethylaminoparthenolide (DMAPT) which lacks a reactive Michael acceptor, impaired multiple components of the FAK1 signaling pathway, including phosphorylation of FAK1 itself, as well as p130Cas and PXN phosphorylation *in situ* in 231MFP breast cancer cells (Fig. 2J-2K). While FAK1 knockdown itself did not affect p130Cas and PXN phosphorylation, FAK1 knockdown conferred total resistance to parthenolide-mediated inhibition of p130Cas and PXN phosphorylation observed in siControl 231MFP cells (Fig. 2L). These results suggest that the slower or longer knockdown of FAK1 by siRNA leads to a rewiring of FAK1 signaling to maintain p130Cas and PXN activity, but that the acute parthenolide-mediated inhibition of FAK1 signaling is still FAK1-dependent and contributes to the viability and motility impairments observed (Fig. 2F-2G).

Activation of FAK1 has also been shown to recruit phosphatidyl inositol-3-kinase (PI3K) to activate AKT/PKC-mediated cell survival pathways (Fig. 2F) (Frame et al., 2010; Sulzmaier et al., 2014). While we observed inhibition of AKT phosphorylation with parthenolide treatment, this inhibition was not attenuated in siFAK1 cells and was thus not mediated by parthenolide interactions with FAK1, but rather through interactions with other targets **(Fig. S1)**. Consistent with known interaction of parthenolide with IKK-βto inhibit IKK-βand NFκB signaling (Kwok et al., 2001), NFκB phosphorylation was inhibited by parthenolide and this inhibition was also not driven through FAK1 **(Fig. S1)**. Nonetheless, our data demonstrates that C427 of FAK1 is both a covalent and functional target of parthenolide that contributes to acute inhibition of specific arms of the FAK1 signaling pathway and the overall anti-cancer effects of parthenolide.

Parthenolide, which belongs to the germacrene family of sesquiterpenes, is just one natural product among hundreds of known sesquiterpene lactones containing the reactive α-methylene-γ-butyrolactone motif (Jackson et al., 2017). Many of these plant metabolites possess notable anti-cancer properties and have been employed in traditional medicine regimes (Ren et al., 2016; Silva Castro et al., 2017). Given the accessibility of C427, we wondered if other related sesquiterpene lactones can also target this residue. The related germacranolide costunolide and the guaianolide natural product dehydrocostus lactone, which both contain α-methylene-γ-butyrolactone pharmacophores, impaired 231MFP breast cancer cell survival (Fig. 3A). On the other hand, the eudesmane-type sesquiterpene natural product α-Santonin, which does not contain this reactive functional group did not impair 231MFP cell survival (Fig. 3A). Consistent with these results, parthenolide, costunolide, and dehydrocostus lactone, but not α-santonin, exhibited FAK1 cysteine reactivity as shown by competitive IA-rhodamine labeling of FAK1 (Fig. 3B). Moreover, the more highly oxidized guaianolide sesquiterpene mikanokryptin (Hu et al., 2017), which also possesses differing stereochemistry relative to dehydrocostus lactone, also showed FAK1 cysteine reactivity and impaired 231MFP breast cancer cell proliferation and survival (Fig. 3C-E). Taken together, these findings indicate some flexibility in targeting this druggable hotspot with sesquiterpene natural products harboring reactive α-methylene-γ-butyrolactone homo-Michael acceptors. We do note, however, that this reactive cysteine has not been pinpointed in previous target identification studies employing related natural products (Tian et al., 2017).

**Figure 3.**
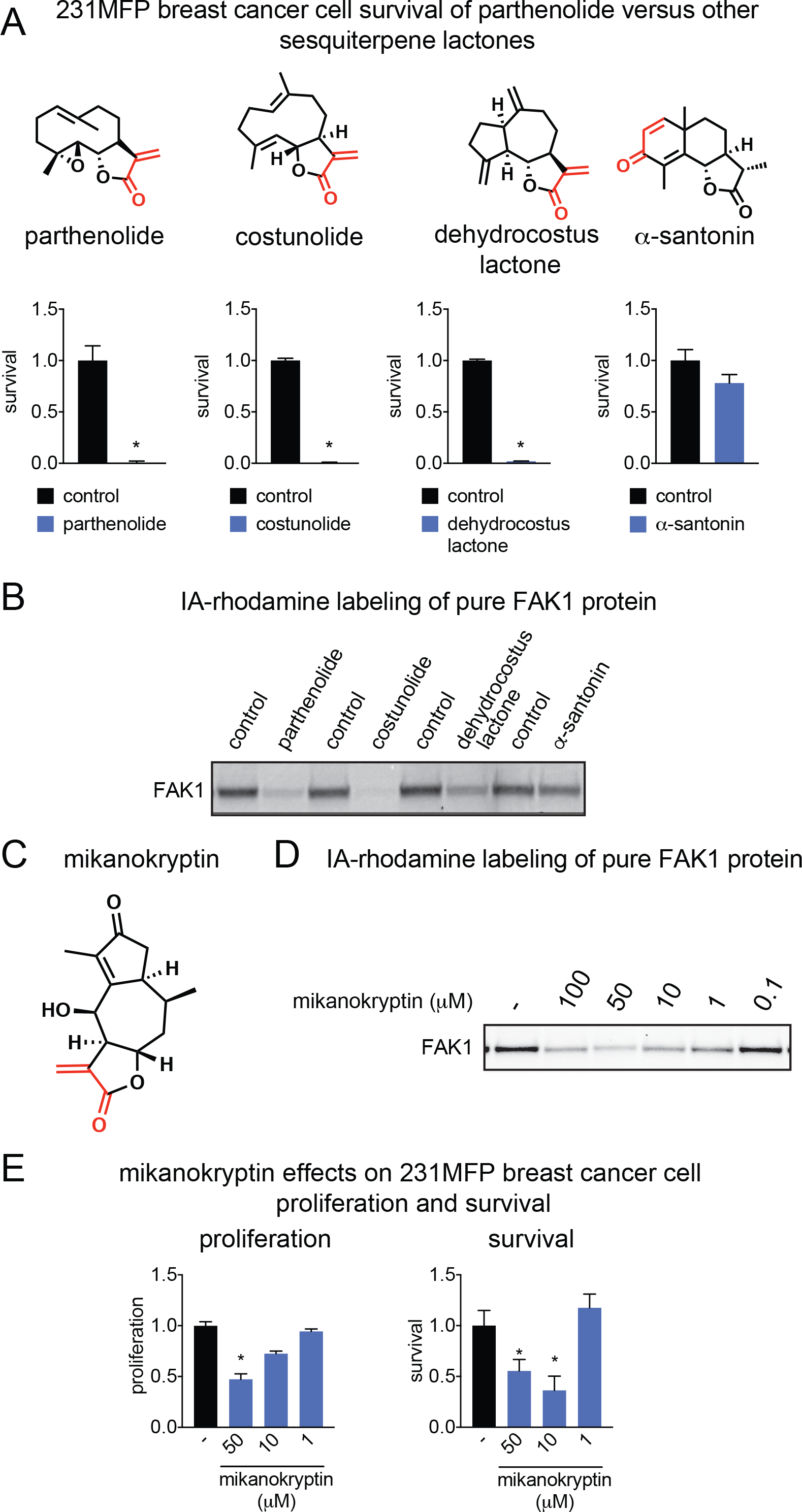
Other sesquiterpene lactones also interact with FAK1 and impair TNBC cell viability. **(A)** 231MFP cell survival (48 h) after treatment with DMSO vehicle or parthenolide and other sesquiterpene lactones (50 μM) assessed by Hoechst stain. **(B)** Gel-based ABPP analysis of parthenolide and other sesquiterpene lactones. Pure human FAK1 kinase domain was pre-treated with DMSO vehicle or natural products (50 μM) for 30 min prior to IA-rhodamine labeling (1 μM) for 1 h and then analyzed by in-gel fluorescence. **(C)** Structure of mikanokryptin. Cysteine-reactive enone highlighted in red. **(D)** Gel-based ABPP analysis of mikanokryptin analyzed as described in **(B)**. **(E)** 231MFP breast cancer cell proliferation and survival (48 h) after treatment of cells with DMSO vehicle or mikanokryptin. Gels shown in **(B, D)** are representative of n=3. Data shown in **(A, E)** are average ± sem, n=6/group. Significance is expressed as *p<0.05 compared to vehicle-treated controls.

In accordance with Chen’s findings (Yen-Pon et al., 2018), we also discovered small, fully-synthetic covalent binders of FAK C427 via a cysteine-reactive covalent ligand screen using gel-based ABPP against the pure human FAK1 kinase domain **(Fig. S2-S4; Table S2).** After screening 149 covalent ligands, we found 15 potential hits that impaired IA-rhodamine labeling of FAK1 **(Fig. S2)**. We then evaluated these 15 compounds for 231MFP survival and proliferation impairment to identify compounds that gave similar responses to parthenolide. This led us to 3 promising leads, namely TRH 1-191, TRH 1-23, and TRH 1-171 **(Fig. S3)**. We then tested these compounds for FAK1 signaling impairment. While all three impaired Y397 FAK1 phosphorylation in 231MFP cells, TRH 1-191 was superior **(Fig. S3)**. We further confirmed that TRH 1-191 impaired FAK1 IA-rhodamine labeling at comparable concentrations to parthenolide without causing any artifactual protein precipitation which may arise from non-specific reactivity **(Fig. S3)**. While TRH 1-191 is a simple chloroacetamide fragment, we showed that a structurally similar negative control compound TRH 1-189 did not react with FAK1 and had no impact on FAK1 signaling **(Fig. S3)**. We also showed the corresponding TRH 1-191 covalent adduct on C427 of FAK1 by LC-MS/MS **(Fig. S4)**. TRH 1-191 also inhibited FAK1 activity *in vitro* with pure FAK1 kinase domain protein and impaired FAK1 signaling *in situ* in 231MFP breast cancer cells **(Fig. S4)**. TRH 1-191 may thus represent an additional scaffold for targeting C427 of FAK1 to inhibit FAK1 signaling in cancer cells.

Here, we have used ABPP-based chemoproteomic platforms to identify C427 of FAK1 as a novel ligandable and functional site targeted by the anti-cancer natural product parthenolide. This target adds further complexity to the polypharmacological landscape of parthenolide which also includes IKK-β, heat shock protein Hsp72, and thioredoxin reductase as targets (Duan et al., 2016; Kwok et al., 2001; Shin et al., 2017). Moreover, these findings also implicate a broad array of widely examined natural products as potential FAK impairment agents–The contribution of these effects to global cytotoxicity warrants further future study among other sesquiterpene lactone natural products. Furthermore, our study highlights the utility of using chemoproteomic platforms for discovering unique druggable modalities that are accessed by covalently-acting natural products.

## Methods

### Chemicals

Parthenolide and dimethylaminoparthenolide were obtained from Cayman Chemicals. Synthesis of the parthenolide-alkyne probe was performed as previously reported (Shin et al., 2017). All other chemicals were obtained from Millipore-Sigma unless otherwise noted. Antibodies were obtained from Cell Signaling Technologies unless otherwise noted. Mikanokryptin was synthesized as previously described (Hu et al., 2017). Synthesis and characterization of cysteine-reactive covalent ligands screened against FAK1 were either described previously or described in Supporting Methods (Bateman et al., 2017; Grossman et al., 2017; Roberts et al., 2017b).

### Cell Culture

Professor Benjamin Cravatt’s group provided the 231MFP cells. The generation of these cells have been described previously (Jessani et al., 2004). HCC38 and HEK293T cells were obtained from American Type Culture Collection (ATCC). 231MFP cells were cultured in L-15 (HyClone) media supplemented with 10% fetal bovine serum (FBS) (Gibco) and 2 mM glutamine at 37° C and 0% CO_2_. HCC38 cells were cultured in RPMI (Gibco) supplemented with 10 % FBS and 2 mM glutamine at 37° C and 5% CO_2_. HEK293T cells were cultured in DMEM (Gibco) supplemented with 10 % FBS and 2 mM glutamine at 37° C and 5% CO_2_.

### Cell proliferation and survival

Serum-containing cell proliferation and serum-free cell survival were assessed by Hoechst stain as previously described (Grossman et al., 2017; Louie et al., 2016). Briefly, we seeded cells at 1 × 10^4^ and 2 × 10^4^ cells/well, respectively, in 150 μL serum-containing or serum-free media in 96-well plates overnight. The next day, cells were treated with an additional 50 μL of DMSO vehicle or compound-containing media for 24 or 48 h before fixation and staining with 10 % formalin and Hoechst 33342 (Invitrogen) according to manufacturer’s protocol. Wells were washed with PBS, and fluorescence was measured using a fluorescent plate reader with an excitation and emission of 350 nm and 461 nm, respectively.

### Measuring cell migration and apoptosis

Cells (5 × 10^4^) were placed in collagen-coated Transwell chambers (Corning) and incubated in DMSO vehicle or compound-containing serum-free media for 6 h. Migrated cells were fixed and stained with Diff-Quik solution (Dade Behring) and non-migrated cells were removed using a cotton swab. Migrated cells were imaged and counted at 200 × magnification. An average of cells in three fields for one migration chamber represents n=1. Apoptotic analyses were performed 2 h following cell exposure (6cm plates, 1 × 10^6^ cells) to DMSO vehicle or compound-containing serum-free media using flow cytometry to measure the percentage of early apoptotic cells (Annexin V positive, propidium iodide negative) and late apoptotic cells (Annexin V positive, propidium iodide positive), as described previously (Anderson et al., 2017). Data analysis was performed using FlowJo software.

### Tumor Xenograft Studies

C.B17 SCID female mice (6-8 weeks old) were injected subcutaneously in the flank with 231MFP cells (1 × 10^6^ cells) suspended in serum-free media. Mice were exposed via intraperitoneal injection with either vehicle (18:1:1 PBS/ethanol/PEG40) or 30 mg/kg parthenolide once per day starting 2 weeks, respectively, after injection of cancer cells. Tumor size was assessed weekly by caliper measurements. Animal experiments were conducted in accordance with the guidelines of the Institutional Animal Care and Use (IACUC) of the University of California, Berkeley.

### RNA interference knockdown of FAK1

Cells were plated in 6-well plates overnight (2 × 10^5^ cells/well). Cells were then treated with Dharmafect1 reagent (GE) and either non-targeting siRNA oligonucleotide (siControl, D-001810-10-05, GE) or siFAK1 oligonucleotides (L-003164-00, GE) were transfected for 48 h according to manufacturer instructions. Cells were then reseeded for survival and proliferation as described above. Knockdown was confirmed by Western blotting.

### Vectors for FAK1 kinase control and C427A overexpression

The full length FAK1 expression vector was purchased from VectorBuilder (FAK1 sequence NM_005607.4). To express just the kinase domain (AA393-698) Gibson Assembly was performed using the pCMV6-Entry (C-term FLAG + Myc tag) using the primers CTGCCGCCGCGATCGCCatggaaacagatgattatgctgagattataga, TCGAGCGGCCGCGTACGCGTtcttctggactccatcctcatgcgctcttcttgct to amplify the FAK1 kinase domain ORF with desired overlaps, and ACGCGTACGCGGCCG, GGCGATCGCGGCGG to linearize the pCMV6-Entry backbone. To achieve the C427A mutant, site-directed mutagenesis was performed using Q5 Site-Directed Mutagenesis Kit (New England BioLabs) using the primers ACTTGGACGAGCTATTGGAGAAGGC, TCTATTCTTTCTCTTTGAATCTC according to manufacturer protocol.

### FAK1 C427A Overexpression

HEK293T cells were seeded at 30% confluency in 15 cm dishes in 10%. On the day of transfection media was replaced with DMEM containing 2.5 % FBS and 500μL Opti-MEM (Thermo) containing 10μg pCMV6-Entry-FAK1 either control or C427A vector and 50 μg polyethylenimine was added to the plate. 48 h later cells were scraped into 1 mL PBS and pelleted at 2,000 g for 5 min at 4° C and the supernatant removed before freezing at −80° C to achieve cell lysis. Pellets were then resuspended in 500 μL PBS and further lysed by probe tip sonication at 15 % amplitude for 2 × 10 s on ice. Lysates were cleared by centrifugation at 21,000 g for 20 min at 4° C and the resulting supernatant was mixed with 30 μL anti-FLAG resin (Genescript) and rotated at 4° C for 2 h before washing 3 × with 500 μL PBS and subsequent elution of FLAG-tagged proteins with 100 μL of 250 ng/μL 3 × FLAG peptide (APExBIO) in PBS. Resultant peptides were further concentrated and 3 × FLAG peptide removed using Amicon centrifugal filtration devices (Millipore).

### Western Blotting

Cells (1 × 10^6^ cells) were plated in 6 cm dishes in complete media the night before experiment. Cells were washed with PBS and placed in DMSO vehicle or compound-containing serum-free media for 2 h before being washed and collected into a lysis buffer containing protease and phosphatase inhibitors. Proteins were separated on a 4-20% Tris-Glycine precast Midi-PROTEAN TGX SDS/PAGE gel (BioRad). Proteins were then transferred to a PVDF membrane using the iBlot system (Invitrogen). Membranes were blocked in 5 % nonfat milk in TBST and incubated in primary antibodies overnight according to manufacturer instructions. Membranes were then washed in TBST and probed with secondary antibody (Li-Cor) and visualized using a fluorescent scanner (Li-Cor). Quantitation was performed using ImageJ.

### FAK1 Activity Assay

FAK1 kinase domain (0.1μg) was preincubated with DMSO vehicle or compound (100 μM) for 30 minutes at room temperature in 5 μL buffer containing 40 mM Tris, 20 mM MgCl_2_, 2 mM MnCl_2_, 4% DMSO, pH 7.4. ATP solution (5 μL of a 100 μM solution containing 0.1 μg 4:1 glycine:tyrosine peptide substrate (Promega)) was added and incubated at 37 °C for 20 min before the addition of ADP-Glo reagent (5μL) (Promega) for an additional 20 min incubation at room temperature. Kinase detection solution (10 μL) (Promega) was added and incubated for 15 minutes at room temperature and luminescence was measured on a plate reader.

### IsoTOP-ABPP analysis of parthenolide targets

Cell lysate was preincubated with DMSO vehicle or parthenolide (50 μM) for 30 minutes at room temperature before labeling of proteomes with iodoacetamide (IA)-alkyne (10 μM) (Chess Organics) for 1 h at room temperature. Isotopically light (control) or heavy (treated) TEV-biotin handles (100 μM) were appended to probe-labeled proteins by CuAAC as previously described. Probe-labeled proteins were subsequently mixed in a 1:1 ratio, avidin-enriched, digested with trypsin, and probe-modified tryptic peptides were subsequently isolated and eluted with TEV protease for subsequent LC-MS/MS analysis as previously described (Grossman et al., 2017; Roberts et al., 2017b).

### Proteomic profiling to determine sites of modification of parthenolide and TRH 1-191

FAK1 kinase domain (AA393-698) (Promega) (25 μg) was pre-incubated with DMSO vehicle or compound (parthenolide or TRH 1-191) (100 μM) for 30 min at room temperature. Samples were then treated with isotopically light (control) or heavy (treated) iodoacetamide (Iodoacetamide-^13^C_2_, 2-d_2_, #721328, Millipore-Sigma) for 1 h at room temperature before combination of control and treated samples and subsequent precipitation with 20% trichloroacetic acid for 2 h at −80° C. Proteins were pelleted at 20,000 × *g* and washed with ice cold 10 mM HCl / 90% acetone before resuspension in 60 μL 4 M Urea and 0.5 × ProteaseMax (Promega), vortexed, and then subsequently diluted with an additional 40 μL 100 mM ammonium bicarbonate. Samples were incubated at 60° C for 30 min following the addition of 10 μL 110 mM TCEP, before dilution with 120 μL 0.04 × ProteaseMax and 5 μg/μL sequencing grade trypsin (Promega) in PBS. Samples were digested overnight at 37° C in a rocking incubator before acidification with 12 μL formic acid and storage at −80° C until MS analysis.

### MS analysis

Total peptides from TEV protease digestion for isoTOP-ABPP or tryptic peptides for shotgun proteomics were pressure loaded onto 250 μm tubing packed with Aqua C18 reverse phase resin (Phenomenex), attached to a nanospray column packed with C18 resin and strong-cation exchange resin, and then analyzed using a Q-Exactive Plus mass spectrometer (Thermo Fisher Scientific) using a Multidimenstional Protein Identification Technology (MudPIT) program as described previously (Grossman et al., 2017; Roberts et al., 2017b).

Following data extraction as reported previously, data were searched against the Uniprot human database using ProLuCID search methodology in IP2 v.3 (Integrated Proteomics Applications, Inc.) (Xu et al., 2015). For isoTOP-ABPP, peptides were analyzed with a static modification for cysteine carboxyaminomethylation (+57.02146) and differential modifications for light or heavy TEV tags (+464.28596 and +470.29977, respectively) for cysteine, and methionine oxidation, with up to two total modifications per peptide. For proteomic analysis of while FAK1 tryptic digests, peptides were searched with differential modifications for light or heavy iodoacetamide (+57.02146 and +61.04073, respectively) on cysteines, for parthenolide or TRH 1-191 addition (+248.14125 or + 259.04001, respectively) on cysteines, and methionine oxidation, with up to two total modifications per peptide fragment. In both analyses, peptides were required to have at least one tryptic end and in isoTOP-ABPP all fragments must contain the TEV modification. To ensure a peptide false-positive of less than 5% ProLuCID data were filtered through DTASelect prior to downstream analyses. Only peptides that were present in at least two out of four biological replicates were interpreted for final quantification and only those peptides with light to heavy ratios >10 that appeared in three out of four biological replicates were considered as targets of parthenolide.

### Gel-Based ABPP

Recombinant FAK1 kinase domain was diluted to .002 μg/μL in PBS and 50 μL protein solution was treated with DMSO vehicle or compound for 30 min at room temperature. Samples were then incubated with IA-rhodmaine (1 μM) (Thermo) for 1 h at room temperature in the dark before separating proteins by SDS/PAGE. Probe-labeled proteins were analyzed by in-gel fluorescence using a Bio-Rad gel scanner and fluorescent bands were quantified using Image J.

## Author Contributions

DKN and CAB conceived of the project, contributed intellectually to the project, and performed, analyzed, and interpreted experiments and data, and wrote the paper; RH, HSL, MT, CRA, SGD performed, analyzed, and interpreted experiments and data; XH, TRH, YP synthesized small-molecules described in this project; TJM, JAO analyzed and interpreted experiments and data, contributed intellectually to the project, and assisted in paper-writing.

## Acknowledgements

We thank the members of the Nomura Research Group for critical reading of the manuscript. This work was supported by the National Institutes of Health (NIH/NCI CA172667, NIH/NIEHS P42ES004705, R01ES028096, and R01GM116952), American Cancer Society Research Scholar Award (RSG-14-242-01-TBE), the Novartis-Berkeley Center for Proteomics and Chemistry Technologies, and the Mark Foundation for Cancer Research ASPIRE award.

