## Supporting Information for "Parthenolide Covalently Targets and Inhibits Focal Adhesion Kinase in Breast Cancer Cells"

### Supporting Methods

#### General synthetic methods

Chemicals and reagents were purchased from major commercial suppliers and used without further purification. Reactions were performed under a nitrogen atmosphere unless otherwise noted. Silica gel flash column chromatography was performed using EMD or Sigma Aldrich silica gel 60 (230-400 mesh). Proton and carbon nuclear magnetic resonance ($^1$H NMR and $^{13}$C NMR) data was acquired on a Bruker AVB 400, AVQ 400, or AV 600 spectrometer at the University of California, Berkeley. High resolution mass spectrum were obtained from the QB3 mass spectrometry facility at the University of California, Berkeley using positive or negative electrospray ionization (+ESI or -ESI). Yields are reported as a single run.

#### General Procedure A

The amine (1 eq.) was dissolved in DCM (5 mL/mmol) and cooled to 0$^o$C. To the solution was added acryloyl chloride (1.2 eq.) followed by triethylamine (1.2 eq.). The solution was warmed to room temperature and stirred overnight. The solution was then washed with brine and the crude product was purified by silica gel chromatography (and recrystallization if necessary) to afford the corresponding acrylamide.

#### General Procedure B

The amine (1 eq.) was dissolved in DCM (5 mL/mmol) and cooled to 0$^o$C. To the solution was added chloroacetyl chloride (1.2 eq.) followed by triethylamine (1.2 eq.). The solution was warmed to room temperature and stirred overnight. The solution was then washed with brine and the crude product was purified by silica gel chromatography (and recrystallization if necessary) to afford the corresponding chloroacetamide.

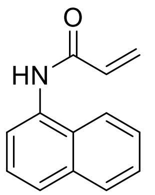


***N*-(naphthalene-1-yl)acrylamide (TRH-1-57).**

To a solution of 1-naphthylamine (294 mg, 2.0 mmol) in dichloromethane (10 mL) was added acryloyl chloride (0.20 mL, 2.4 mmol) followed by triethylamine (248 mg, 2.4 mmol) at 0˚ C under $N_2$ atmosphere. The reaction mixture was allowed to warm to room temperature and was stirred for 16 hours. The solution was washed twice with brine, and the resulting crude was purified by silica gel chromatography (30% to 40% ethyl acetate in hexanes) and recrystallized from toluene to yield 173 mg of white solid (44% yield).

**$^1$H NMR (400MHz, MeOD)**: δ 7.96-7.94 (m, 1H), 7.88-7.86 (m, 1H), 7.76 (d, *J* = 8.1 Hz, 1H), 7.64 (d, *J* = 7.3 Hz, 1H), 7.52-7.44 (m, 3H), 6.43 (dd, *J* = 16.9, 10.4 Hz, 1H), 6.41 (dd, *J* = 16.9, 1.5 Hz, 1H), 5.82 (dd, *J* = 10.1, 1.0 Hz, 1H).

**$^{13}$C NMR (100MHz, MeOD)**: δ 167.3, 135.7, 133.9, 132.1, 129.9, 129.4, 128.2, 127.7, 127.3, 127.2, 126.4, 123.8, 123.3.

**HRMS (+ESI)**: Calculated: 198.0913 ($C_{13}H_{12}NO$). Observed: 198.0912.

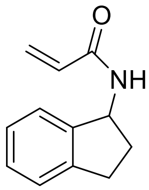

**N-(2,3-dihydro-1*H*-inden-1-yl)acrylamide (TRH-1-58).**
To a solution of 1-aminoindan (274 mg, 2.0 mmol) in dichloromethane (10 mL) was added acryloyl chloride (0.20 mL, 2.4 mmol) followed by triethylamine (276 mg, 2.4 mmol) at 0˚ C under $N_2$ atmosphere. After stirring for 20 minutes, the reaction mixture was allowed to warm to room temperature and was stirred for 28 hours. The solution was washed twice with brine, and the resulting crude was purified by silica gel chromatography (20%-40% ethyl acetate in hexanes) to yield 238 mg of white solid (62% yield).
**$^{1}$H NMR (400MHz, CDCl$_3$)**: δ 7.22-7.11 (m, 4H), 6.76 (d, *J* = 7.8 Hz, 1H), 6.23-6.13 (m, 2H), 5.56 (dd, *J* = 3.5, 8.1 Hz, 1H), 5.40 (q, *J* = 7.9 Hz, 1H), 2.95-2.88 (m, 1H), 2.84-2.76 (m, 1H), 2.51-2.43 (m, 1H), 1.83-1.74 (m, 1H).
**$^{13}$C NMR (100MHz, CDCl$_3$)**: δ 165.5, 143.2, 143.1, 130.9, 127.8,126.6, 126.3, 124.6, 124.0, 54.5, 33.7, 30.2.
**HRMS (+ESI)**: Calculated: 188.1070 ($C_{12}H_{14}NO$). Observed: 188.1068.

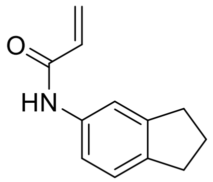

***N*-(2,3-dihydro-1*H*-inden-5-yl)acrylamide (TRH-1-59).**
To a solution of 5-aminoindan (269 mg, 2.0 mmol) in dichloromethane (10 mL) was added acryloyl chloride (0.20 mL, 2.4 mmol) followed by triethylamine (270 mg, 2.4 mmol) at 0˚ C under $N_2$ atmosphere. The reaction mixture was allowed to warm to room temperature and was stirred for 20 hours. The solution was washed twice with brine, and the resulting crude was purified by silica gel chromatography (40% ethyl acetate in hexanes) to yield 129 mg of white solid (34% yield).
**$^{1}$H NMR (400MHz, CDCl$_3$)**: δ 8.87 (s, 1H), 7.54 (s, 1H), 7.31 (d, *J* = 7.6 Hz, 1H), 7.11 (d, *J* = 7.8 Hz, 1H), 6.40 (d, *J* = 5.6 Hz, 2H), 5.66 (t, *J* = 5.6 Hz, 1H), 2.87-2.80 (m, 4H), 2.04 (t, *J* = 7.2 Hz, 2H).
**$^{13}$C NMR (100MHz, CDCl$_3$)**: δ 164.4, 144.9, 140.4, 136.0, 131.6, 127.0, 124.3, 118.8, 117.0, 32.9, 32.4, 25.6.
**HRMS (+ESI)**: Calculated: 188.1070 ($C_{12}H_{14}NO$). Observed: 188.1068.

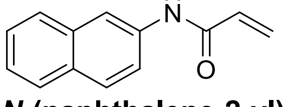

***N*-(naphthalene-2-yl)acrylamide (TRH-1-60).**
To a solution of 2-naphthylamine (289 mg, 2.0 mmol) in dichloromethane (10 mL) was added acryloyl chloride (0.20 mL, 2.4 mmol) followed by triethylamine (269 mg, 2.4 mmol) at 0˚ C under $N_2$ atmosphere. After 15 minutes, the reaction mixture was allowed to warm to room temperature and was stirred for 16 hours. The solution was washed twice with 5% citric acid and once with brine, and the resulting crude was purified by

silica gel chromatography (30% ethyl acetate in hexanes) to yield 266 mg of an off-white solid (67% yield).
**$^1$H NMR (400MHz, MeOD)**: δ 8.25 (d, *J* = 1.8 Hz, 1H), 7.73-7.69 (m, 3H), 5.54 (dd, *J* = 2.1, 8.8 Hz, 1H), 7.39-7.34 (m, 1H), 7.33-7.29 (m, 1H), 6.44 (dd, *J* = 9.7, 17.0 Hz, 1H), 6.36 (dd, *J* = 2.2, 17.0 Hz, 1H), 5.72 (dd, *J* = 2.2, 9.7 Hz, 1H).
**$^{13}$C NMR (100MHz, MeOD)**: δ 166.2, 137.1, 135.1, 132.4, 132.1, 129.5, 128.6, 128.5, 127.9, 127.4, 126.1, 121.1, 118.1.
**HRMS (+ESI)**: Calculated: 198.0913 ($C_{13}H_{12}NO$). Observed: 198.0912.

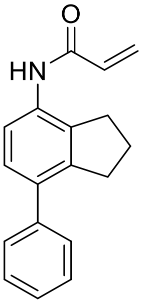


**N-(7-phenyl-2,3-dihydro-1*H*-inden-4-yl)acrylamide (TRH-1-68).**

To a solution of *N*-(7-bromo-2,3-dihydro-1*H*-inden-4yl)acrylamide (**TRH-1-65**, 56 mg, 0.2 mmol) in a solution of dioxane and water (4:1 dioxane:water, 2.1 mL) was added sequentially phenylboronic acid (55 mg, 0.4 mmol), potassium carbonate (78 mg, 0.5 mmol), and tetrakis(triphenylphosphine)palladium(0) (26 mg, 10 mol%). The reaction mixture was heated to a reflux and was stirred overnight. The reaction was diluted with water (20 mL) and extracted with DCM (3x20 mL). The combined organics were evaporated and the resulting crude was purified by silica gel chromatography (10% to 50% ethyl acetate in hexanes) then recrystallized from toluene to give 11 mg of white solid (20% yield).

**$^1$H NMR (400MHz, MeOD)**: δ 7.48 (d, *J* = 8.2 Hz, 1H), 7.40-7.39 (m, 4H), 7.33-7.27 (m, 1H), 7.16 (d, *J* = 8.2 Hz, 1H), 6.53 (dd, *J* = 10.2, 17.0 Hz, 1H), 6.36 (dd, *J* = 1.7, 17.0 Hz, 1H), 5.77 (dd, *J* =1.7, 10.2 Hz, 1H), 2.96 (t, *J*= 7.3 Hz, 2H), 2.91 (t, *J* = 7.3 Hz, 2H), 2.04 (quint, *J* = 7.3 Hz, 2H).

**$^{13}$C NMR (100MHz, MeOD)**: δ 166.3, 144.2, 142.4, 139.2, 133.8, 132.2, 129., 129.3, 128.2, 127.9, 127.8, 123.0, 34.3, 32.1, 26.6.

**HRMS (+ESI)**: Calculated: 264.1383 ($C_{18}H_{18}NO$). Observed: 264.1381.

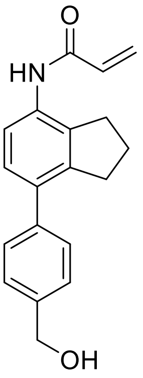


***N*-(7-(4-(hydroxymethyl)phenyl)-2,3-dihydro-1*H*-4-yl)acrylamide (TRH-1-70).**

To a solution of *N*-(7-bromo-2,3-dihydro-1*H*-inden-4yl)acrylamide (**TRH-1-65**, 56 mg, 0.2 mmol) in a solution of dioxane and water (4:1 dioxane:water, 2.1 mL) under nitrogen atmosphere was added sequentially 4-(hydroxymethyl)phenylboronic acid (66 mg, 0.4 mmol), potassium carbonate (78 mg, 0.5 mmol), and tetrakis(triphenylphosphine)palladium(0) (26 mg, 10 mol%). The reaction mixture was heated to a reflux and stirred overnight. The reaction was diluted with water (20 mL) and extracted with DCM (3x20 mL). The combined organics were dried with magnesium sulfate, filtered, and evaporated, and the resulting crude was purified by silica gel chromatography (20% to 50% ethyl acetate in hexanes) to give 16 mg of white solid (26% yield).

**$^{1}$H NMR (400MHz, MeOD)**: δ 7.47 (d, *J* = 8.1 Hz, 1H), 7.41-7.38 (m, 4H), 7.15 (d, *J* = 8.2 Hz, 1H), 6.52 (dd, *J* = 10.2, 16.9 Hz, 1H), 6.36 (d, *J* = 17.0 Hz, 1H), 5.77 (d, *J* = 10.5 Hz, 1H), 4.63 (s, 2H), 2.95 (t, *J* = 7.1 Hz, 2H), 2.90 (t, *J* = 7.2 Hz, 2H), 2.03 (t, *J* = 7.3 Hz, 2H).

**$^{13}$C NMR (100MHz, MeOD)**: δ 166.3, 144.1, 141.4, 139.2, 136.9, 133.8, 132.2, 129.5, 128.5,128.2, 128.0, 127.9, 123.0, 65.0, 34.3, 32.0, 26.6.

**HRMS (+ESI)**: Calculated: 294.1489 ($C_{19}H_{20}NO_2$). Observed: 294.1486.

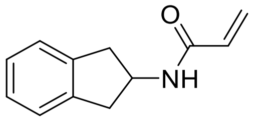


***N*-(2,3-dihydro-1*H*-inden-2-yl)acrylamide (TRH-1-74).**

To a solution of 2-aminoindan (253 mg, 2.0 mmol) in dichloromethane (10 mL) was added acryloyl chloride (0.19 mL, 2.4 mmol) followed by triethylamine (222 mg, 2.4 mmol) at 0˚ C under $N_2$ atmosphere. The reaction mixture was allowed to warm to room temperature after 15 minutes and was stirred for 23 hours. The solution was washed twice with brine, and the resulting crude was purified by silica gel chromatography (40% ethyl acetate in hexanes) to yield 126 mg of an off-white solid (35% yield).

**$^1$H NMR (400MHz, $CDCl_3$)**: δ 7.23-7.16 (m, 4H), 6.27 (dd, *J* = 1.3, 17.0 Hz, 1H), 6.10 (s, 1H), 6.04 (dd, *J* = 10.3, 17.0 Hz, 1H), 5.60 (dd, *J* = 1.3, 10.3 Hz, 1H), 4.82-4.75 (m, 1H), 3.32 (dd, *J* = 7.1, 16.2 Hz, 2H), 2.84 (dd, *J* = 4.4, 16.1 Hz, 2H).

**$^{13}$C NMR (100MHz, $CDCl_3$)**: δ 165.4 ,140.9, 130.9, 126.8, 126.5, 124.9, 50.7, 40.1.

**HRMS (+ESI)**: Calculated: 188.1070 ($C_{12}H_{14}NO$). Observed: 188.1068.

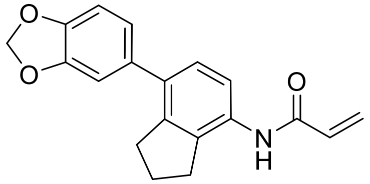


***N*-(7-(benzo[*d*][1,3]dioxol-5-yl)-2,3-dihydro-1*H*-inden-4-yl)acrylamide (TRH-1-78).**
To a solution of N-(7-bromo-2,3-dihydro-1H-inden-4yl)acrylamide (**TRH-1-65**, 55 mg, 0.2 mmol) in a mixture of dioxane and water (4:1 dioxane:water, 2.1 mL) under nitrogen atmosphere was added sequentially 3,4-(methylenedioxy)phenylboronic acid (70 mg, 0.4 mmol), potassium carbonate (74 mg, 0.5 mmol), and tetrakis(triphenylphosphine)palladium(0) (24 mg, 10 mol%). The reaction mixture was heated to a reflux and stirred overnight. The reaction was diluted with water (20 mL) and extracted with DCM (3x20 mL). The combined organics were dried with magnesium sulfate, filtered, and evaporated, and the resulting crude was purified by silica gel chromatography (0% to 25% ethyl acetate in hexanes) to give 7 mg of white solid (11% yield).

**$^1$H NMR (600 MHz, $CDCl_3$)**: δ 7.93 (d, *J* = 7.0 Hz, 1H), 7.18 (d, *J* = 8.2 Hz, 1H), 7.10 (s, 1H), 6.90 (s, 1H), 6.86 (t, *J* = 8.1 Hz, 2H), 6.45 (d, *J* = 16.8 Hz, 1H), 6.30 (dd, *J* = 10.3, 16.8 Hz, 1H), 5.79 (d, *J* = 10.2 Hz, 1H), 3.00 (t, *J* = 7.3 Hz, 2H), 2.88 (t, *J* = 7.3 Hz, 2H), 2.10 (quint, 7.3 Hz, 2H).
**$^{13}$C NMR (150 MHz, $CDCl_3$)**: δ 147.7, 146.7, 142.84, 142.81, 135.2, 134.9, 132.8, 131.3, 127.9, 127.8, 122.1, 119.8, 109.2, 108.3, 101.2, 36.8, 33.5, 30.5, 25.4.
**HRMS (-ESI)**: Calculated: 306.1136 ($C_{19}H_{16}NO_3$). Observed: 306.1130.

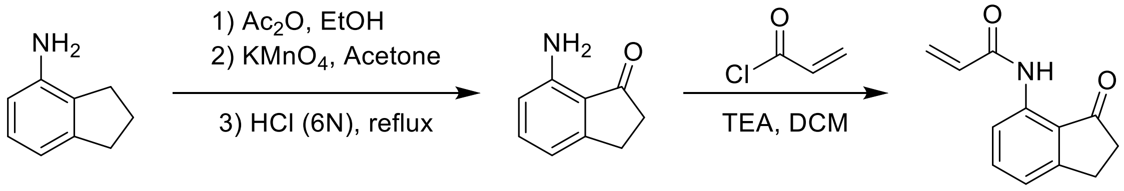


***N*-(3-oxo-2,3-dihydro-1*H*-inden-4-yl)acrylamide (TRH-1-129)**
To a solution of 4-aminoindan (1.0 g, 7.5 mmol) in ethanol (20 mL) at 0 °C was added acetic anhydride (1.4 mL, 15.0 mmol). The solution was raised to room temperature and stirred overnight, after which the solvent was evaporated. The residue was then dissolved in acetone (50 mL) to which was added 15% aqueous magnesium sulfate (1.2 g in 6.75 mL of water) followed by potassium permanganate (3.4 g, 17.0 mmol), and the resulting solution was stirred for 24 hours. The reaction filtered through a pad of celite, eluting with chloroform and then water. The eluent was separated, and the aqueous layer was extracted several times with additional chloroform. The combined organics were dried over magnesium sulfate, filtered and evaporated. The residue was then dissolved in a 6N HCl solution (20 mL) and heated to 90 °C. After stirring for 5 hours, the solution was cooled, neutralized with small portions of potassium carbonate, and

extracted with ethyl acetate. The combined organics were dried with magnesium sulfate, filtered, and evaporated to give 610 mg (55% over 3 steps) of crude **7-aminoindan-1-one** which was used without further purification.

To a solution of 7-aminoindan-1-one in dichloromethane (15 mL) was added acryloyl chloride (0.39 mL, 4.8 mmol) followed by triethylamine (0.67 mL, 4.8 mmol) at 0˚ C under $N_2$ atmosphere. The reaction mixture was allowed to warm to room temperature and was stirred overnight. The solution was washed 1M HCl solution (2x) and brine, and the resulting crude was purified by silica gel chromatography (10% to 20% ethyl acetate in hexanes) to yield 390 mg of white solid (47% yield, 26% combined over 4 steps).

**$^1$H NMR (400MHz, $CDCl_3$)**: δ 10.64 (s, 1H), 8.45 (d, *J* = 8.2 Hz, 1H), 7.55 (t, *J* = 7.9 Hz, 1H), 7.12 (d, *J* = 7.6 Hz, 1H), 6.45 (dd, *J* = 1.0, 17.0 Hz, 1H), 6.33 (dd, *J* = 10.1, 17.0 Hz, 1H), 5.82 (dd, *J* = 1.0, 10.1 Hz, 1H), 3.11 (t, *J* = 11.5 Hz, 2H), 2.74-2.71 (m, 2H).

**$^{13}$C NMR (100MHz, $CDCl_3$)**: δ 209.3, 164.4, 155.9, 138.7, 137.0, 131.7, 128.0, 123.1, 120.8, 116.9, 36.5, 25.5.

**HRMS (+ESI)**: Calculated: 202.0863 ($C_{12}H_{12}NO_2$). Observed: 202.0860.

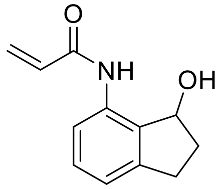


***N*-(3-hydroxy-2,3-dihydro-1*H*-inden-4-yl)acrylamide (TRH-1-133).**

To a solution of *N*-(3-oxo-2,3-dihydro-1*H*-inden-4-yl)acrylamide (**TRH-1-129**, 201 mg, 1.0 mmol) in anhydrous methanol (7 mL) under nitrogen atmosphere was added sodium borohydride (46.1 mg, 1.2 mmol). After 30 minutes of stirring, the reaction was quenched with saturated sodium bicarbonate solution and extracted three times with DCM. The combined organics were dried with magnesium sulfate, filtered, and concentrated. Crude was purified by silica gel chromatography (30 to 50% ethyl acetate in hexanes) to give 190 mg of the product as a white solid (94% yield).

**$^1$H NMR (400 MHz, $CDCl_3$)**: δ 8.93 (s, 1H), 7.98 (d, *J* = 7.8 Hz, 1H), 7.19 (t, *J* = 7.9 Hz, 1H), 6.95 (d, *J* = 7.4 Hz, 1H), 6.29 (d, *J* = 16.8 Hz, 1H), 6.15 (dd, *J* = 10.2, 16.9 Hz, 1H), 5.66 (d, *J* = 10.2 Hz, 1H), 5.32 (q, *J* = 6.9 Hz, 1H), 3.60 (d, *J* = 6.7 Hz, 1H), 2.96 (ddd, *J* = 2.4, 9.0, 15.7 Hz), 2.73 (quint, *J* = 8.1 Hz, 1H), 2.56-2.48 (m, 1H), 1.96-1.86 (m, 1H).

**$^{13}$C NMR (100 MHz, $CDCl_3$)**: δ 164.1, 143,7 ,135.6, 132.8, 131.6, 129.5, 127.3, 121.0, 118.5, 76.2, 36.0, 29.8.

**HRMS (-ESI)**: Calculated: 202.0874 ($C_{12}H_{12}NO_2$). Observed: 202.0874.

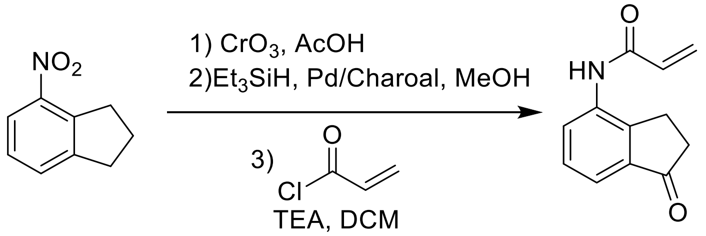


***N*-(1-oxo-2,3-dihydro-1*H*-inden-4-yl)acrylamide (TRH-1-134).**

To a solution of 4-nitroindan (5.38 g, 33 mmol) in acetic acid (250 mL) was slowly added chromium trioxide (8.95 g, 90 mmol). After stirring for 24 hours, the reaction was neutralized with 2M NaOH and extracted five times with ethyl acetate. The combined organics were washed with a saturated sodium bicarbonate solution and brine, then

dried over magnesium sulfate, filtered, and concentrated. The crude material was purified by silica gel chromatography (10-20% ethyl acetate in hexanes) to give 1.26 g (ca. 7.1 mmol) of 4-nitroindanone as a white solid.
This intermediate was combined with palladium on activated charcoal (125 mg, 10 wt%) dissolved in anhydrous methanol (21 mL) under the atmosphere of a nitrogen balloon. Triethylsilane (11.3 mL, 71 mmol) was slowly added by addition funnel over the course of 10 minutes while the reaction was stirred under the cooling of a room temperature water bath. After an additional 20 minutes of stirring, the reaction mixture was filtered through a pad of celite and subsequently concentrated to give crude 4-aminoindanone which was used without further purification.
This final intermediate was then dissolved in DCM (21 mL) under $N_2$ atmosphere and cooled to 0°C, after which acryloyl chloride (0.77 mL, 9.5 mmol) and triethylamine (1.19 mL, 8.5 mmol) were slowly added. The reaction was allowed to warm to room temperature while stirring overnight, at which point the reaction was washed twice with brine, dried with magnesium sulfate, filtered, and concentrated. The crude was purified by silica gel chromatography (30-50% ethyl acetate in hexanes) to give 989 mg of a white solid (15% yield over 3 steps).
**$^1$H NMR (400 MHz, $CDCl_3$)**: δ 8.20 (d, *J* = 5.8 Hz, 1H), 7.63 (s, 1H), 7.56 (d, *J* = 7.5 Hz, 1H), 7.39 (t, *J* = 7.7 Hz, 1H), 6.48 (d, *J* = 16.7 Hz, 1H), 6.37 (dd, *J* = 10.0 Hz, 16.8 Hz, 1H), 5.83 (d, *J* = 10.1 Hz, 1H), 3.04 (t, *J* = 5.6 Hz, 2H), 2.70 (t, *J* = 5.7 Hz, 2H).
**$^{13}$C NMR (100 MHz, $CDCl_3$)**: δ 206.3, 163.9, 146.0, 138.0, 135.4, 130.7, 128.8, 128.7, 127.6, 120.4, 36.1, 23.4.
**HRMS (-ESI)**: Calculated: 200.0717 ($C_{12}H_{10}NO_2$). Observed: 200.0715.

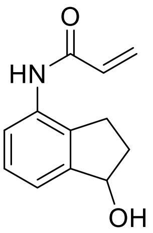


***N*-(1-hydroxy-2,3-dihydro-1*H*-inden-4-yl)acrylamide (TRH-1-135).**
To a solution of *N*-(1-oxo-2,3-dihydro-1*H*-inden-4-yl)acrylamide (**TRH-1-134**, 1.26 g, 6.25 mmol) in anhydrous methanol (50 mL) under nitrogen atmosphere was added sodium borohydride (292.7 mg, 7.7 mmol). After 30 minutes of stirring, the reaction was quenched with water and the methanol was removed *in vacuo*. The residue was saturated with NaCl and extracted five times with a 2:1 chloroform:methanol solution. The combined organics were dried over 3 angstrom molecular sieves, filtered, and concentrated. The crude material was purified by silica gel chromatography (40 to 70% ethyl acetate in hexanes) to give 1.05 g of the product as a white solid (83% yield).
**$^1$H NMR (400 MHz, MeOD)**: δ 7.50 (dd, *J* = 2.3, 6.3 Hz, 1H), 7.25-7.20 (m, 2H), 6.51 (dd, *J* = 10.2, 17.0 Hz, 1H), 6.35 (dd, *J* = 1.7, 17.0 Hz, 1H), 5.77 (dd, *J* = 1.7, 10.2 Hz, 1H), 5.17 (t, *J* = 6.3 Hz, 1H), 2.97 (ddd, *J* = 4.5, 8.6, 16.2, 1H), 2.74 (quint, *J* = 7.8 Hz, 1H), 2.47-2.39 (m, 1H), 1.95-1.86 (m, 1H).
**$^{13}$C NMR (100 MHz, MeOD)**: δ 166.3, 148.0, 137.8, 134.8, 132.1, 128.3, 127.9, 124.0, 122.7, 76.9, 36.1, 28.6.
**HRMS (-ESI)**: Calculated: 202.0874 ($C_{12}H_{12}NO_2$). Observed: 202.0872.

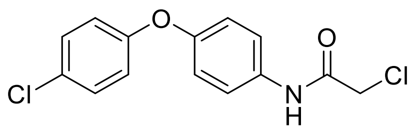

**2-Chloro-*N*-(4-(4-chlorophenoxy)phenyl)acetamide (TRH-1-140).**
To a solution 4-(4-chlorophenoxy)aniline (446 mg, 2.0 mmol) in dichloromethane (10 mL) was added chloroacetyl chloride (0.39 mL, 4.8 mmol) followed by triethylamine (0.67 mL, 4.8 mmol) at 0˚ C under $N_2$ atmosphere. After stirring for 35 minutes, the reaction mixture was allowed to warm to room temperature and was stirred for 27 hours. The solution was washed twice with brine, dried with magnesium sulfate, and the resulting crude was purified by silica gel chromatography (30% to 50% ethyl acetate in hexanes) to yield 533 mg of an off-white solid (89% yield).
**$^1$H NMR (400 MHz, $CDCl_3$)**: δ 8.33 (s, 1H), 7.52-7.48 (m, 2H), 7.29-7.25 (m, 2H), 6.99-6.96 (m, 2H), 6.93-6.89 (m, 2H), 4.17 (s, 2H)
**$^{13}$C NMR (100 MHz, $CDCl_3$)**: δ 164.1, 156.0, 154.0, 132.5, 129.8, 128.4, 122.2, 119.9, 119.6, 42.9.
**HRMS (-ESI)**: Calculated: 294.0094 ($C_{14}H_{10}NO_2Cl_2$). Observed: 294.0094.

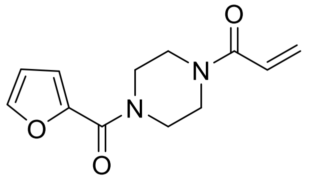

**1-(4-(furan-2-carbonyl)piperazin-1-yl)prop-2-en-1-one (TRH-1-145).**
To a solution 1-(2-furoyl)piperazine (362 mg, 2.0 mmol) in dichloromethane (10 mL) was added acryloyl chloride (0.20 mL, 2.4 mmol) followed by triethylamine (0.34 mL, 2.4 mmol) at 0˚ C under $N_2$ atmosphere. After stirring for 20 minutes, the reaction mixture was allowed to warm to room temperature and was stirred for 24 hours. The solution was washed twice with brine, dried with magnesium sulfate, and the resulting crude was purified by silica gel chromatography (70% to 100% ethyl acetate in hexanes) to yield 446 mg of yellow solid (95% yield).
**$^1$H NMR (400 MHz, $CDCl_3$)**: δ 7.53 (m, 1H), 7.06 (dd, *J* = 0.7, 3.5 Hz, 1H), 6.61 (dd, *J* = 10.5, 16.8 Hz, 1H), 6.52 (dd, *J* = 1.8, 3.5 Hz, 1H), 6.33 (dd, *J* = 1.9, 16.8 Hz, 1H), 5.75 (dd, *J* = 1.9, 10.5 Hz, 1H), 3.84-3.67 (m, 8H).
**$^{13}$C NMR (100 MHz, $CDCl_3$)**: δ 165.5, 159.1, 147.5, 144.0, 128.5, 127.1, 117.0, 111.5, 45.6, 41.9.
**HRMS (+ESI)**: Calculated: 235.1077 ($C_{12}H_{15}N_2O_3$). Observed: 235.1075.

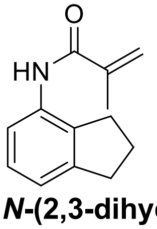

***N*-(2,3-dihydro-1*H*-inden-4-yl)methacrylamide (TRH-1-149).**
To a solution 4-aminoindan (0.24 mL, 2.0 mmol) in dichloromethane (10 mL) was added methacryloyl chloride (0.23 mL, 2.4 mmol) followed by triethylamine (0.34 mL, 2.4 mmol) at 0˚ C under $N_2$ atmosphere. After stirring for 20 minutes, the reaction mixture was allowed to warm to room temperature and was stirred for 3.5 hours. The solution was washed twice with brine, dried with magnesium sulfate, and the resulting crude was

purified by silica gel chromatography (35% to 40% ethyl acetate in hexanes) to yield 378 mg of off-white solid (94% yield).

**$^{1}$H NMR (400 MHz, CDCl$_3$)**: δ 7.72 (d, *J* = 8.0 Hz, 1H), 7.55 (s, 1H), 7.12 (t, *J* = 7.7 Hz, 1H), 7.01 (d, *J* = 7.4 Hz, 1H), 5.79 (s, 1H), 5.42 (s, 1H), 2.93 (t, *J* = 7.5 Hz, 2H), 2.79 (t, *J* =7.4 Hz, 2H), 7.12-2.06 (m, 2H), 2.04 (s, 3H).

**$^{13}$C NMR (100 MHz, CDCl$_3$)**: δ 166.3, 145.1, 140.6, 134.5, 133.7, 127.0, 120.7, 119.8, 118.9, 33.1, 29.9, 24.7, 18.6.

**HRMS (+ESI)**: Calculated: 202.1226 ($C_{13}H_{16}NO$). Observed: 202.1224.

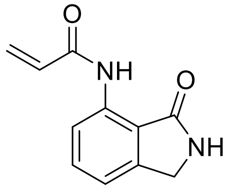


***N*-(3-oxoisoindolin-4-yl)acrylamide (TRH-1-152).**

To a solution of 7-aminoisoindolin-1-one (99 mg, 0.67 mmol) in dichloromethane (4 mL) was added acryloyl chloride (0.07 mL, 0.8 mmol) followed by triethylamine (0.11 mL, 0.8 mmol) at 0˚ C under $N_2$ atmosphere. After stirring for 20 minutes, the reaction mixture was allowed to warm to room temperature and was stirred overnight. The solution was washed twice with brine, dried with magnesium sulfate, and the resulting crude was purified by silica gel chromatography (50 to 60% ethyl acetate in hexanes) to yield 58 mg of a white solid (43% yield).

**$^{1}$H NMR (400 MHz, CDCl$_3$)**: δ 10.50 (s, 1H), 8.58 (d, *J* = 8.2 Hz, 1H), 7.55 (t, *J* = 7.9 Hz, 1H), 7.15 (d, *J* = 7.5 Hz, 1H), 6.82 (s, 1H), 6.46 (dd, *J* = 1.3, 17.0 Hz, 1H), 6.36 (dd, *J* = 10.0, 17.0 Hz, 1H), 5.81 (dd, *J* = 1.3, 10.0 Hz, 1H), 4.46 (s, 2H).

**$^{13}$C NMR (100 MHz, CDCl$_3$)**: δ 172.9, 164.2, 143.9, 138.2, 133.8, 131.8, 127.8, 118.0, 117.7, 117.6, 45.6.

**HRMS (+ESI)**: Calculated: 203.0815 ($C_{11}H_{11}N_2O_2$). Observed: 203.0814.

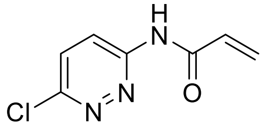


***N*-(6-chloropyridazin-3-yl)acrylamide (TRH-1-155).**

To a solution 3-amino-6-chloropyridazine (261 mg, 2.0 mmol) in dichloromethane (10 mL) was added acryloyl chloride (0.20 mL, 2.4 mmol) followed by triethylamine (0.34 mL, 2.4 mmol) at 0˚ C under $N_2$ atmosphere. After stirring for 20 minutes, the reaction mixture was allowed to warm to room temperature and was stirred overnight. The solution was washed twice with brine, dried with magnesium sulfate, and the resulting crude was purified by silica gel chromatography (40% to 50% ethyl acetate in hexanes) to yield 23 mg of a pale-yellow solid (6% yield).

**$^{1}$H NMR (400 MHz, CDCl$_3$)**: δ 10.06 (s, 1H), 8.70 (d, *J* = 9.4 Hz, 1H), 7.57 (d, *J* = 9.4 Hz, 1H), 6.73 (dd, *J* = 10.2, 16.8 Hz, 1H) 6.56 (dd, *J* = 1.2, 16.8, 1H), 5.94 (dd, *J* = 1.2, 10.2 Hz, 1H).

**$^{13}$C NMR (100 MHz, CDCl$_3$)**: δ 164.8, 155.2, 152.3, 130.7, 130.4, 130.3, 122.0.

**HRMS (+ESI)**: Calculated: 182.0127 ($C_7H_5N_3OCl$). Observed: 182.0126.

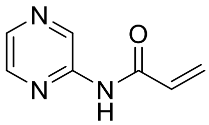

***N*-(Pyrazin-2-yl)acrylamide (TRH-1-156).**
To a solution of aminopyrazine (192 mg, 2.0 mmol) in dichloromethane (10 mL) was added acryloyl chloride (0.20 mL, 2.4 mmol) followed by triethylamine (0.34 mL, 2.4 mmol) at 0˚ C under $N_2$ atmosphere. After stirring for 20 minutes, the reaction mixture was allowed to warm to room temperature and was stirred overnight. The solution was washed twice with brine, dried with magnesium sulfate, and the resulting crude was purified by silica gel chromatography (50% to 70% ethyl acetate in hexanes) to yield 22 mg of white solid (7% yield).
**$^1$H NMR (600 MHz, $CDCl_3$)**: δ 9.65 (d, *J* = 1.3 Hz, 1H), 8.38 (d, *J* = 2.5 Hz, 1H), 8.27 (dd, *J* = 1.6, 2.5 Hz, 1H), 8.19 (s, 1H), 6.54 (dd, *J* = 0.8, 16.9 Hz, 1H), 6.33 (dd, *J* = 10.3, 16.9 Hz, 1H), 5.90 (dd, *J* = 0.8, 10.3 Hz, 1H).
**$^{13}$C NMR (150 MHz, $CDCl_3$)**: δ 163.5, 148.2, 142.2, 140.6, 137.4, 130.2, 129.8.
**HRMS (+ESI)**: Calculated: 150.0662 ($C_7H_8N_3O$). Observed: 150.0660.

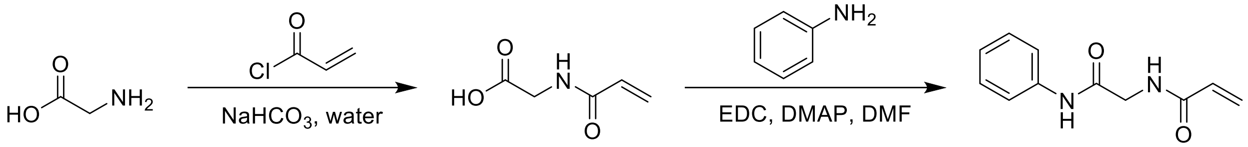


***N*-(2-oxo-2-(phenylamino)ethyl)acrylamide (TRH-1-160).**
To a solution of glycine (1.50 g, 20.0 mmol) and sodium bicarbonate (1.70 g, 20.2 mmol) in water (30 mL) at 0˚ C was slowly added acryloyl chloride (2.45 mL, 30.2 mmol). After stirring for 3.5 hours, the reaction was extracted 3 times with ethyl acetate. The combined organics were dried over magnesium sulfate, filtered, and concentrated to give an oil. The oil was treated with hexanes causing a white solid to crash out which was collected by gravity filtration to give 124 mg of crude acryloylglycine of which 58 mg (47% of the crude material) was used immediately without further purification.
This solid (ca. 0.45 mmol) was dissolved in DMF (2.5 mL) and a solution of N-(3-dimethylaminopropyl)-N′-ethylcarbodiimide hydrochloride (104 mg, 0.54 mmol) and 4-dimethylaminopyridine (68 mg, 0.56 mmol) in DMF (2.5 mL) was added followed by aniline (0.050 mL, 0.54 mmol). The solution was stirred overnight, diluted with ethyl acetate, and washed with both a saturated solution of sodium bicarbonate and brine. The organics were then dried over magnesium sulfate, filtered, and concentrated, and the resulting crude was purified by silica gel chromatography (30-60% ethyl acetate in hexanes) to give 19 mg of the title compound as a white solid (1% yield over two steps).

**$^1$H NMR (400 MHz, MeOD)**: δ 7.56-7.53 (m, 2H), 7.30 (t, *J* = 8.0 Hz, 2H), 7.08 (t, *J* = 7.4 Hz, 1H), 6.35 (dd, *J* = 9.9, 17.1 Hz, 1H), 6.26 (dd, *J* = 2.0, 17.1 Hz, 1H), 5.71 (dd, *J* = 2.0, 9.9 Hz, 1H), 4.08 (s, 2H).
**$^{13}$C NMR (100 MHz, MeOD)**: δ 169.4, 168.6, 139.5, 131.7, 129.8, 127.2, 125.3, 121.2, 44.0.
**HRMS (-ESI)**: Calculated: 203.0826 ($C_{11}H_{11}N_2O_2$). Observed: 203.0825.

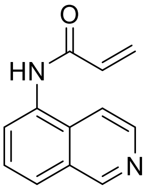

***N*-(isoquinolin-5-yl)acrylamide (TRH-1-162).**

To a solution of 5-aminoisoquinoline (287 mg, 2.0 mmol) in dichloromethane (10 mL) was added acryloyl chloride (0.20 mL, 2.4 mmol) followed by triethylamine (0.34 mL, 2.4 mmol) at 0˚ C under $N_2$ atmosphere. After stirring for 20 minutes, the reaction mixture was allowed to warm to room temperature and was stirred overnight. The solution was washed with brine, and the resulting aqueous layer was extracted with a 2:1 chloroform:methanol solution. The resulting crude was purified by chromatography on basic alumina (50% ethyl acetate in hexanes to 4% ethanol in ethyl acetate) to yield 43 mg of a yellow solid (11% yield).

**$^1$H NMR (400 MHz, MeOD)**: δ 9.23 (s, 1H), 8.45 (d, *J* = 6.1 Hz, 1H), 8.03 (d, *J* = 7.5 Hz, 1H), 7.97 (d, *J* = 8.2 Hz, 1H), 7.88 (d, *J* = 6.1 Hz, 1H), 7.68 (t, *J* = 7.9 Hz, 1H), 6.66 (dd, *J* = 10.2, 17.0 Hz, 1H), 6.47 (dd, *J* = 1.5, 17.0 Hz, 1H), 5.88 (dd, *J* = 1.7, 10.2 Hz, 1H).

**$^{13}$C NMR (100 MHz, MeOD)**: δ 167.1, 153.5, 143.0, 133.6, 132.4, 131.9, 130.6, 128.7, 127.9, 127.1, 117.2.

**HRMS (+ESI)**: Calculated: 199.0866 ($C_{12}H_{11}N_2O$). Observed: 199.0863.

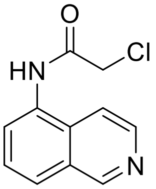

**2-Chloro-*N*-(isoquinolin-5-yl)acetamide (TRH-1-163).**

To a solution 5-aminoisoquinoline (289 mg, 2.0 mmol) in dichloromethane (10 mL) was added chloroacetyl chloride (0.19 mL, 2.4 mmol) followed by triethylamine (0.34 mL, 2.4 mmol) at 0˚ C under $N_2$ atmosphere. After stirring for 20 minutes, the reaction mixture was allowed to warm to room temperature and was stirred overnight. The solution was washed with saturated sodium bicarbonate solution and brine, dried with magnesium sulfate, and the resulting crude was purified by chromatography on basic alumina (30% ethyl acetate in hexanes to 4% ethanol in ethyl acetate) to yield 157 mg of yellow solid (36% yield).

**$^1$H NMR (600 MHz, MeOD)**: δ 9.26 (s, 1H), 8.48 (d, *J* = 6.1 Hz, 1H), 8.03 (d, *J* = 8.2 Hz, 1H), 7.98 (d, *J* = 7.4 Hz, 1H), 7.91 (d, *J* = 6.1 Hz, 1H), 7.71 (t, *J* = 7.9 Hz, 1H), 4.39, (s, 2H).

**$^{13}$C NMR (150 MHz, MeOD)**: δ 167.4, 152.1, 141.7, 131.8, 131.2, 129.2, 127.3, 127.0, 126.1, 115.7, 42.3.

**HRMS (+ESI)**: Calculated: 221.0476 ($C_{11}H_{10}N_2O$). Observed: 221.0473.

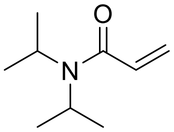

***N,N*-diisopropylacrylamide (TRH-1-167).**

To a solution of diisopropylamine (0.42 mL, 3.0 mmol) in dichloromethane (10 mL) was added acryloyl chloride (0.29 mL, 3.6 mmol) followed by triethylamine (0.50 mL, 3.6

mmol) at 0˚ C under $N_2$ atmosphere. After stirring for 20 minutes, the reaction mixture was allowed to warm to room temperature and was stirred for 19 hours. The solution was washed with a saturated solution of sodium bicarbonate followed by brine, dried with magnesium sulfate, and the resulting crude was purified by silica gel chromatography (0% to 30% ethyl acetate in hexanes) to yield 392 mg of a pale-yellow oil (84% yield).
**$^1$H NMR (400 MHz, $CDCl_3$)**: δ 6.35 (dd, *J* = 10.6, 16.8 Hz, 1H), 5.98 (dd, *J* = 1.7, 16.8 Hz, 1H), 5.36 (dd, *J* = 1.7, 10.6 Hz, 1H), 3.85 (s, 1H), 3.56 (s, 1H), 1.18 (s, 6H), 1.06 (s, 6H).
**$^{13}$C NMR (100 MHz, $CDCl_3$)**: δ 165.9, 130.5, 125.3, 47.9, 45.4, 21.1, 20.3.
**HRMS (+ESI)**: Calculated: 178.1202 ($C_9H_{17}NONa$). Observed: 178.1201.

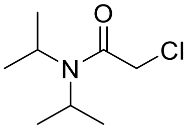

**2-Chloro-*N,N*-diisopropylacetamide (TRH-1-168).**
To a solution diisopropylamine (0.42 mL, 3.0 mmol) in dichloromethane (10 mL) was added chloroacetyl chloride (0.29 mL, 3.6 mmol) followed by triethylamine (0.50 mL, 3.6 mmol) at 0˚ C under $N_2$ atmosphere. After stirring for 20 minutes, the reaction mixture was allowed to warm to room temperature and was stirred overnight. The solution was washed with saturated sodium bicarbonate solution and brine, dried with magnesium sulfate, and the resulting crude was purified by silica gel chromatography (0 to 20% ethyl acetate in hexanes) to yield 376 mg of white solid (70% yield).
**$^1$H NMR (400 MHz, $CDCl_3$)**: δ 3.93 (s, 2H), 3.88-3.82 (m, 1H), 3.38-3.31 (m, 1H), 1.29 (d, *J* = 6.5 Hz, 6H), 1.14 (d, *J* = 6.4 Hz, 6H).
**$^{13}$C NMR (100 MHz, $CDCl_3$)**: δ 165.0, 49.7, 46.1, 43.2, 20.7, 20.0.
**HRMS (+ESI)**: Calculated: 200.0813 ($C_8H_{16}NOClNa$). Observed: 200.0811.

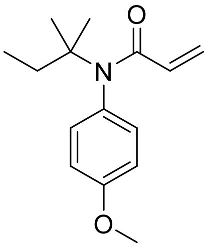

***N*-(4-methoxyphenyl)-*N*-(*tert*-pentyl)acrylamide (TRH-1-170).**
To a solution of 4-methoxy-*N*-(*tert*-pentyl)aniline (94 mg, 0.49 mmol) in dichloromethane (5 mL) was added acryloyl chloride (0.05 mL, 0.6 mmol) followed by triethylamine (0.09 mL, 0.6 mmol) at 0˚ C under $N_2$ atmosphere. After stirring for 15 minutes, the reaction mixture was allowed to warm to room temperature and was stirred for 18 hours. The solution was washed with a saturated solution of sodium bicarbonate followed by brine, dried with magnesium sulfate, and the resulting crude was purified by silica gel chromatography (0% to 20% ethyl acetate in hexanes) to yield 82 mg of a pale-yellow oil (68% yield).
**$^1$H NMR (400 MHz, $CDCl_3$)**: δ 6.99 (d, *J* = 8.7 Hz, 2H), 6.85 (d, *J* = 8.7 Hz, 2H), 6.17 (dd, *J* = 1.9, 16.7 Hz, 1H), 5.76 (dd, *J* = 10.3, 16.7 Hz, 1H), 5.28 (dd, *J* = 1.9, 10.3 Hz, 1H), 3.81 (s, 3H), 2.11 (q, *J* = 7.5 Hz, 2H), 1.20 (s, 6H), 0.91 (t, *J* = 7.5 Hz, 3H).
**$^{13}$C NMR (100 MHz, $CDCl_3$)**: δ 166.3, 159.0, 134.3, 131.49, 131.45, 125.6, 114.1, 61.7, 55.5, 32.0, 27.4, 9.4.

**HRMS (+EI)**: Calculated: 247.1572 ($C_{15}H_{21}NO_2$). Observed: 247.1577.

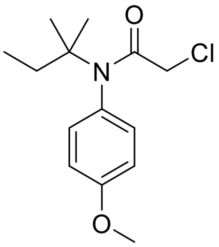


**2-Chloro-*N*-(4-methoxyphenyl)-*N*-(*tert*-pentyl)acetamide (TRH-1-171).**
To a solution 4-methoxy-*N*-(*tert*-pentyl)aniline (95 mg, 0.5 mmol) in dichloromethane (5 mL) was added chloroacetyl chloride (0.05 mL, 0.6 mmol) followed by triethylamine (0.085 mL, 0.6 mmol) at 0˚ C under $N_2$ atmosphere. After stirring for 15 minutes, the reaction mixture was allowed to warm to room temperature and was stirred overnight. The solution was washed with saturated sodium bicarbonate solution and brine, dried with magnesium sulfate, and the resulting crude was purified by silica gel chromatography (0 to 10% ethyl acetate in hexanes) to yield 99 mg of a yellow oil (74% yield).
**$^1$H NMR (400 MHz, $CDCl_3$)**: δ 7.04 (d, *J* = 8.6 Hz, 2H), 6.87 (d, *J* = 8.6 Hz, 2H), 3.80 (s, 3H), 3.63 (s, 2H), 2.05 (q, *J* = 7.4 Hz, 2H), 1.16 (s, 6H), 0.90 (t, *J* = 7.4 Hz, 3H).
**$^{13}$C NMR (100 MHz, $CDCl_3$)**: δ 166.0, 159.5, 133.2, 131.0, 114.5, 62.5, 55.5, 44.8, 31.8, 27.1, 9.3.
**HRMS (+ESI)**: Calculated: 270.1255 ($C_{14}H_{21}NO_2Cl$). Observed: 270.1254.

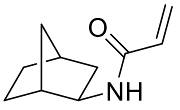


***N*-(*exo*-norborn-2-yl)acrylamide (TRH-1-176).**
To a solution of *exo*-2-aminonorbornane (0.24 mL, 2 mmol) in dichloromethane (10 mL) was added acryloyl chloride (0.20 mL, 2.4 mmol) followed by triethylamine (0.33 mL, 2.4 mmol) at 0˚ C under $N_2$ atmosphere. After stirring for 20 minutes, the reaction mixture was allowed to warm to room temperature and was stirred for 18 hours. The solution was washed with a saturated solution of sodium bicarbonate followed by brine, dried with magnesium sulfate, and the resulting crude was purified by silica gel chromatography (30% ethyl acetate in hexanes) to yield 271 mg of a white solid (82% yield).
**$^1$H NMR (400 MHz, $CDCl_3$)**: δ 6.42 (s, 1H), 6.25 (dd, *J* = 2.3, 17.0 Hz, 1H), 6.18 (dd, *J* = 9.5, 17.0 Hz, 1H), 5.58 (dd, *J* = 2.3, 9.5 Hz, 1H), 3.8-3.77 (m, 1H), 2.27-2.24 (m, 2H), 1.78 (ddd, *J* = 2.1, 8.1, 13.0 Hz, 1H), 1.55-1.38 (m, 3H), 1.30-1.10 (m, 4H).
**$^{13}$C NMR (100 MHz, $CDCl_3$)**: δ 165.0, 131.4, 125.8, 52.9, 42.4, 40.0, 35.7, 35.6, 28.2, 26.6.
**HRMS (+EI)**: Calculated:165.1154 ($C_{10}H_{15}NO$). Observed: 165.1155.

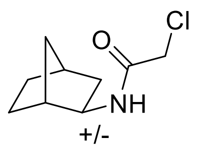


**2-Chloro-*N*-(*exo*-norborn-2-yl)acetamide (TRH-1-177).**
To a solution of *exo*-2-aminonorbornane (0.24 mL, 2 mmol) in dichloromethane (10 mL) was added chloroacetyl chloride (0.19 mL, 2.4 mmol) followed by triethylamine (0.33 mL, 2.4 mmol) at 0˚ C under $N_2$ atmosphere. After stirring for 20 minutes, the reaction mixture was allowed to warm to room temperature and was stirred overnight. The solution was washed with saturated sodium bicarbonate solution and brine, dried with magnesium sulfate, and the resulting crude was purified by silica gel chromatography (20 to 40% ethyl acetate in hexanes) to yield 345 mg of a white solid (91% yield).
**$^1$H NMR (400 MHz, $CDCl_3$)**: δ 6.48 (s, 1H), 3.93 (s, 2H), 3.67-3.63 (m, 1H), 2.24-2.22 (m, 1H), 2.16-2.15 (m, 1H), 1.74 (ddd, *J* = 1.9, 8.1, 13.0 Hz, 1H), 1.50-1.36 (m, 2H), 1.30-1.26 (m, 1H), 1.21-1.14 (m, 3H), 1.09-1.03 (m, 1H).
**$^{13}$C NMR (100 MHz, $CDCl_3$)**: δ 165.0, 53.1, 42.6, 42.2, 40.0, 35.6, 35.5, 28.0, 26.3.
**HRMS (+ESI)**: Calculated: 187.0764 ($C_9H_{14}NOCl$). Observed: 187.0765.

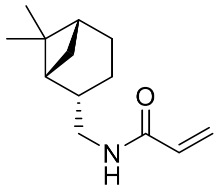


***N*-(((1*R*,2*S*,5*R*)-6,6-dimethylbicyclo[3.1.1]heptan-2-yl)methyl)acrylamide (TRH-1-178).**
To a solution of (-)-*cis*-myrtanylamine (0.34 mL, 2 mmol) in dichloromethane (10 mL) was added acryloyl chloride (0.20 mL, 2.4 mmol) followed by triethylamine (0.33 mL, 2.4 mmol) at 0˚ C under $N_2$ atmosphere. After stirring for 20 minutes, the reaction mixture was allowed to warm to room temperature and was stirred for 21 hours. The solution was washed with a saturated solution of sodium bicarbonate followed by brine, dried with magnesium sulfate, and the resulting crude was purified by silica gel chromatography (20 to 30% ethyl acetate in hexanes) to yield 369 mg of a white solid (89% yield).
**$^1$H NMR (600 MHz, $CDCl_3$)**: δ 6.26 (dd, *J* = 1.5, 17.0 Hz, 1H), 6.11 (dd, *J* = 10.3, 17.0 Hz, 1H) 5.85 (s, 1H), 5.61 (dd, *J* = 1.5, 10.3 Hz, 1H), 3.39-3.29 (m, 2H), 2.38-2.34 (m, 1H), 2.26-2.21 (m, 1H), 1.98-1.90 (m, 4H), 1.88-1.83 (m, 1H), 1.53-1.47 (m, 1H), 1.19 (s, 3H), 1.04 (s, 3H), 0.89 (d, *J* = 9.6 Hz, 1H).
**$^{13}$C NMR (150 MHz, $CDCl_3$)**: δ 165.7, 131.2, 126.2, 45.3, 43.9, 41.5, 38.8, 33.3, 28.1, 26.1, 23.3, 19.9.
**HRMS (-ESI)**: Calculated: 206.1550 ($C_{13}H_{20}NO$). Observed: 206.1551.

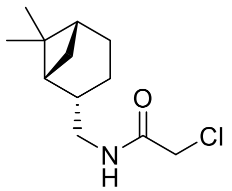

**2-Chloro-*N*-(((1*R*,2*S*,5*R*)-6,6-dimethylbicyclo[3.1.1]heptan-2-yl)methyl)acetamide (TRH-1-179).**
To a solution of (-)-*cis*-myrtanylamine (0.34 mL, 2 mmol) in dichloromethane (10 mL) was added chloroacetyl chloride (0.19 mL, 2.4 mmol) followed by triethylamine (0.33 mL, 2.4 mmol) at 0˚ C under $N_2$ atmosphere. After stirring for 20 minutes, the reaction mixture was allowed to warm to room temperature and was stirred overnight. The solution was washed with saturated sodium bicarbonate solution and brine, dried with magnesium sulfate, and the resulting crude was purified by silica gel chromatography (0 to 20% ethyl acetate in hexanes) to yield 405 mg of an off-white solid (88% yield).
**$^1$H NMR (600 MHz, $CDCl_3$)**: δ 6.61 (s, 1H), 4.05 (s, 2H), 3.33-3.30 (m, 2H), 2.40-2.36 (m, 1H), 2.27-2.21 (m, 1H), 1.99-1.83 (m, 5H), 1.53-1.46 (m, 1H), 1.20 (s, 3H), 1.05 (s, 3H), 0.90 (d, *J* = 9.7 Hz, 1H).
**$^{13}$C NMR (150 MHz, $CDCl_3$)**: δ 165.8, 45.5, 43.8, 42.9, 41.4, 41.2, 38.8, 33.3, 28.0, 26.0, 23.3, 19.8.
**HRMS (-ESI)**: Calculated: 228.1161 ($C_{12}H_{19}NOCl$). Observed: 228.1162.

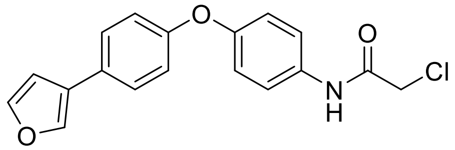

**2-Chloro-*N*-(4-(4-(furan-3-yl)phenoxy)phenyl)acetamide (TRH-1-189).**
A reaction vial equipped with a stirbar was charged with 2-chloro-*N*-(4-(4-chlorophenoxy)phenyl)acetamide (**TRH-1-140**, 74 mg, 0.25 mmol), 3-furanylboronic acid (44 mg, 0.38 mmol), and XPhos-G3-palladacycle (4 mg, 2 mol%) and placed under a nitrogen atmosphere. THF (1 mL) and an aqueous solution of tribasic potassium phosphate (0.5M, 2 mL) that was freshly degassed by sparging with $N_2$ were sequentially added, and the reaction was stirred for 1 hour. The reaction mixture was diluted with water and extracted three times with diethyl ether. The combined organics were dried over magnesium sulfate, and the resulting crude was purified by silica gel chromatography (10% to 30% ethyl acetate in hexanes) to give 19 mg of a white solid (23% yield).
**$^1$H NMR (400 MHz, $CDCl_3$)**: δ 7.50-7.47 (m, 2H), 7.43 (dt, *J* = 2.6, 9.8 Hz, 2H), 7.30-7.24 (m, 3H), 6.95 (dt, *J* = 2.6, 9.7 Hz, 2H), 6.90 (dt, *J* = 2.7, 9.7 Hz, 2H), 6.43 (s, 1H), 3.58 (s, 2H).
**$^{13}$C NMR (100 MHz, $CDCl_3$)**: δ 168.5, 156.2, 153.3, 144.2, 141.2, 133.4, 129.7, 128.1, 121.8, 119.7, 119.6, 117.9, 111.2, 33.9.
**HRMS (-ESI)**: Calculated: 326.0589 ($C_{18}H_{13}NO_3Cl$). Observed: 326.0594.

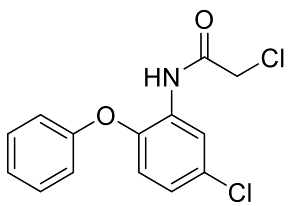


**2-Chloro-*N*-(5-chloro-2-phenoxyphenyl)acetamide (TRH-1-191).**
To a solution of 2-amino-4-chlorophenyl phenyl ether (2.20 g, 10.0 mmol) in dichloromethane (30 mL) was added chloroacetyl chloride (0.96 mL, 12.0 mmol) followed by triethylamine (1.67 mL, 12 mmol) at 0˚ C under $N_2$ atmosphere. After stirring for 20 minutes, the reaction mixture was allowed to warm to room temperature and was stirred for 16 hours. The solution was diluted with DCM, washed with a saturated sodium bicarbonate solution and brine, dried with magnesium sulfate, and the resulting crude was recrystallized from hexanes to yield 1.84 g of light-brown solid (62% yield).
**$^{1}$H NMR (400 MHz, $CDCl_3$)**: δ 8.93 (s, 1H), 8.50 (d, *J* = 2.5 Hz, 1H), 7.40-7.35 (m, 2H), 7.19-7.15 (m, 1H), 7.04-7.01 (m, 3H), 6.81 (d, *J* = 8.7 Hz, 1H), 4.16 (s, 2H).
**$^{13}$C NMR (100 MHz, $CDCl_3$)**: δ 164.0, 156.0, 144.7, 130.2, 129.6, 129.2, 124.7, 124.5, 120.6, 118.71, 118.68, 43.1.
**HRMS (-ESI)**: Calculated: 294.0094 ($C_{14}H_{10}NO_2Cl_2$). Observed: 294.0101.

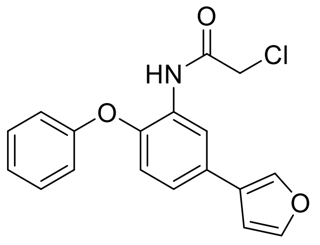


**2-Chloro-*N*-(5-(furan-3-yl)-2-phenoxyphenyl)acetamide (TRH-1-194).**
A reaction vial equipped with a stirbar was charged with 2-chloro-*N*-(5-chloro-2-phenoxyphenyl)acetamide (**TRH-1-191**, 297 mg, 1.0 mmol), 3-furanylboronic acid (171 mg, 1.5 mmol), and XPhos-G3-palladacycle (17 mg, 2 mol%) and placed under a nitrogen atmosphere. THF (2 mL) and a freshly degassed aqueous solution of tribasic potassium phosphate (0.5M, 4 mL) were sequentially added, and the reaction was stirred for 6 hours. The reaction mixture was diluted with water and extracted three times with diethyl ether. The combined organics were dried over magnesium sulfate, and the resulting crude was purified by silica gel chromatography (10% to 15% ethyl acetate in hexanes) to give 120 mg of a yellow solid (37% yield).
**$^{1}$H NMR (400 MHz, $CDCl_3$)**: δ 8.53 (d, *J* = 2.4 Hz, 1H), 7.93 (s, 1H), 7.33-7.25 (m, 4H), 7.12 (t, *J* = 7.4 Hz, 1H), 6.97 (dd, *J* = 2.5, 8.7 Hz, 1H), 6.84 (d, *J* = 7.8 Hz, 2H), 6.79 (d, *J* = 8.7 Hz, 1H), 6.23 (s, 6.23), 3.52 (s, 2H).
**$^{13}$C NMR (100 MHz, $CDCl_3$)**: δ 168.6, 156.3, 144.0, 143.4, 141.1, 130.9, 130.1, 129.7, 124.0, 123.9, 120.7, 119.6, 117.5, 117.4, 111.0, 34.2.
**HRMS (-ESI)**: Calculated: 326.0589 ($C_{18}H_{13}NO_3Cl$). Observed: 326.0585.

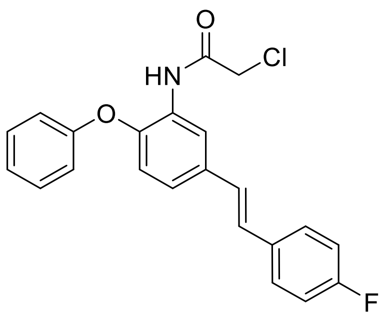

***trans*-2-Chloro-*N*-(5-(4-fluorostyryl)-2-phenoxyphenyl)acetamide (TRH-1-196).**
A reaction vial equipped with a stirbar was charged with 2-chloro-*N*-(5-chloro-2-phenoxyphenyl)acetamide (**TRH-1-191**, 296 mg, 1.0 mmol), *trans*-2-(4-fluorophenyl)vinylboronic acid (253 mg, 1.5 mmol), and XPhos-G3-palladacycle (17 mg, 2 mol%) and placed under a nitrogen atmosphere. THF (2 mL) and a freshly degassed aqueous solution of tribasic potassium phosphate (0.5M, 4 mL) were sequentially added, and the reaction was stirred for 21 hours. The reaction mixture was diluted with water and extracted three times with diethyl ether. The combined organics were dried over magnesium sulfate, and the resulting crude was purified by silica gel chromatography (5% to 15% ethyl acetate in hexanes) to give 256 mg of a yellow solid (67% yield).
**$^{1}$H NMR (400 MHz, $CDCl_3$)**: δ 8.54 (d, *J* = 2.4 Hz, 1H), 8.00 (s, 1H), 7.27-7.20 (m, 4H), 7.10-7.06 (m, 1H), 6.99-6.93 (m, 3H), 6.84-6.79 (m, 3H), 6.47 (d, *J* = 15.9 Hz, 1H), 6.17-6.10 (m, 1H), 3.27 (dd, *J* = 1.1, 7.3 Hz, 2H).
**$^{13}$C NMR (100 MHz, $CDCl_3$)**: δ 168.8, 163.8, 161.3, 156.2, 143.5, 134.6, 132.59, 132.56, 130.9, 130.1, 129.6, 128.1, 128.0, 123.98, 123.96, 121.20, 121.18, 120.7, 119.5, 117.6, 115.7, 115.4, 41.8.
**HRMS (+ESI)**: Calculated: 404.0824 ($C_{22}H_{17}NO_2ClFNa$). Observed: 404.0828.

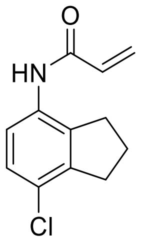

**N-(7-chloro-2,3-dihydro-1H-inden-4-yl)acrylamide (YP-1-1)**
A solution of N-(2,3-dihydro-1H-inden-4-yl)acrylamide (187 mg, 1.0 mmol) in PEG 400 (5.2 mL) was cooled to 0 °C. To the solution was added *N*-chlorosuccinimide (140 mg, 1.0 mmol). The solution was allowed to warm to room temperature after 30 min and stirred overnight. The solution was diluted with ethyl acetate and washed two times with brine and dried with magnesium sulfate. The crude product was purified via silica gel chromatography (30% ethyl acetate in hexanes). The obtained mixture of isomers was separated by recrystallization to afford the product in 22% yield as a white solid (47 mg).
**$^{1}$H NMR (400MHz, $CDCl_3$)**: δ 7.78 (d, *J* = 8.8 Hz, 1H), 7.15-7.11 (m, 2H), 6.42 (dd, *J* = 1.4, 16.8 Hz, 1H), 6.26 (dd, *J* = 10.2, 16.8 Hz, 1H), 5.77 (dd, *J* = 1.4, 10.2 Hz, 1H), 2.98 (t, *J* = 7.6 Hz, 2H), 2.87 (t, *J* = 7.5 Hz, 2H), 2.12 (quint, *J* = 7.5 Hz, 2 H).
**$^{13}$C NMR (100MHz, $CDCl_3$)**: δ 163.4, 143.1, 136.1, 132.2, 131.0, 128.0, 127.2, 126.7, 120.9, 32.7, 31.1, 24.0.
**HRMS (+ESI)**: Calculated: 220.0535 ($C_{12}H_{11}ClNO$). Observed: 220.0533.

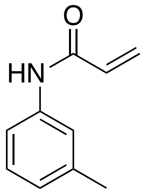

**N-(m-tolyl)acrylamide (YP-1-16)**
A solution of o-toluidine (107 mg, 1.0 mmol) in DCM (10 mL) was cooled to 0 °C. To the solution was added acryloyl chloride (109 mg, 1.2 mmol) followed by triethylamine (121 mg, 1.2 mmol). The solution was allowed to warm to room temperature after 40 min and stirred overnight. The solution was washed two times with brine and dried with magnesium sulfate. The crude product was purified via silica gel chromatography (20% to 40% ethyl acetate in hexanes) to afford the product in 86% yield as a white solid (139 mg).
**$^1$H NMR (400MHz, CDCl$_3$)**: δ 7.82 (d, *J* = 7.9 Hz, 1H), 7.32 (s, 1H), 7.21-7.17 (m, 2H), 7.10-7.06 (m, 1H), 6.43-6.38 (m, 1H), 6.29 (dd, *J* = 10.2, 17.1 Hz, 1H), 5.75-5.72 (m, 1H), 2.25 (s, 1H).
**$^{13}$C NMR (100MHz, CDCl$_3$)**: δ 135.5, 131.2, 130.5, 127.5, 126.8, 125.5, 123.4, 17.8.
**HRMS (+ESI)**: Calculated: 162.0913 ($C_{10}H_{12}NO$). Observed: 162.0912.

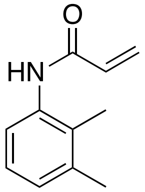

**N-(2,3-dimethylphenyl)acrylamide (YP-1-18)**
A solution of 2,3-dimethylaniline (121 mg, 1.0 mmol) in DCM (10 mL) was cooled to 0 °C. To the solution was added acryloyl chloride (109 mg, 1.2 mmol) followed by triethylamine (121 mg, 1.2 mmol). The solution was allowed to warm to room temperature after 29 min and stirred overnight. The solution was washed two times with brine and dried with magnesium sulfate. The crude product was purified via silica gel chromatography (30% to 40% ethyl acetate in hexanes) to afford the product in 88% yield as a white solid (154 mg).
**$^1$H NMR (400MHz, CDCl$_3$)**: δ 7.49 (d, *J* = 7.9 Hz, 1H), 7.29 (s, 1H), 7.11-7.07 (m, 1H), 7.01 (d, *J* = 7.7, 1H) 6.40 (d, *J* = 17.1, 1H), 6.30 (dd, *J* = 7.3, 17.1 Hz, 1H), 5.74 (d, *J* = 10.1 Hz, 1H), 2.29 (s, 1H), 2.13 (s, 1H).
**$^{13}$C NMR (100MHz, CDCl$_3$)**: δ 135.1, 131.2, 127.6, 127.3, 125.9, 122.3, 20.6, 13.9.
**HRMS (+ESI)**: Calculated: 176.1070 ($C_{11}H_{14}NO$). Observed: 176.1068.

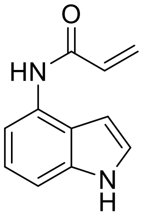

**N-(1H-indol-4-yl)acrylamide (YP-1-19)**
A solution of 4-aminoindole (132 mg, 1 mmol) in DCM (5 mL) and DMF (5 mL) was cooled to 0 °C. To the solution was added acryloyl chloride (109 mg, 1.2 mmol) followed

by triethylamine (121 mg, 1.2 mmol). The solution was allowed to warm to room temperature after 26 min and stirred overnight. The solution was washed two times with brine and dried with magnesium sulfate. The crude product was purified via basic alumina chromatography (60% to 75% ethyl acetate in hexanes) to afford the product in 30% yield as a white-grey solid (56 mg).
**$^{1}$H NMR (600MHz, MeOD)**: δ 7.51 (d, *J* = 7.6 Hz, 1H), 7.24-7.22 (m, 2H), 7.08 (t, *J* = 7.6 Hz, 1H), 6.64 (dd, *J* = 10.1, 16.7 Hz, 2H), 6.38 (dd, *J* = 1.7, 16.9 Hz, 1H), 5.78 (dd, *J* = 1.7, 10.3 Hz, 1H), 4.6 (s, 1H).
**$^{13}$C NMR (150MHz, MeOD)**: δ 165.0, 137.2, 131.1, 129.2, 126.0, 123.8, 121.5, 120.9, 112.2, 108.4, 98.5.
**HRMS (+ESI)**: Calculated: 187.0866 ($C_{11}H_{11}N_2O$). Observed: 187.0865.

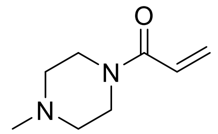

**1-(4-Methylpiperazin-1-yl)prop-2-en-1-one (YP-1-22)**
A solution of 1-methylpiperazine (100 mg, 1.0 mmol) in DCM (10 mL) was cooled to 0 °C. To the solution was added acryloyl chloride (109 mg, 1.2 mmol) followed by triethylamine (121 mg, 1.2 mmol). The solution was allowed to warm to room temperature after 30 min and stirred overnight. The solution was washed two times with brine and dried with magnesium sulfate. The crude product was purified via silica gel chromatography (85% to 100% ethyl acetate in hexanes) to afford the product in 29% yield as a yellow gel (44 mg).
**$^{1}$H NMR (400MHz, $CDCl_3$)**: δ 6.56 (dd, *J* = 10.6, 16.9 Hz, 1H), 6.29 (dd, *J* = 2.0, 16.8 Hz, 1H), 5.69 (dd, *J* = 2.0, 10.6 Hz, 1H), 3.71 (s, 2H), 3.58 (s, 2H), 2.42 (t, *J* = 5.1 Hz, 4H), 2.32 (s, 3H).
**$^{13}$C NMR (100MHz, $CDCl_3$)**: δ 165.4, 127.8, 127.5, 55.2, 54.6, 46.0, 45.7, 41.9.
**HRMS (+ESI)**: Calculated: 155.1179 ($C_8H_{15}N_2O$). Observed: 155.1178.

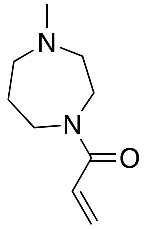

**1-(4-methyl-1,4-diazepan-1-yl)prop-2-en-1-one (YP-1-23)**
A solution of 1-methylhomopiperazine (114 mg, 1.0 mmol) in DCM (10 mL) was cooled to 0 °C. To the solution was added acryloyl chloride (109 mg, 1.2 mmol) followed by triethylamine (121 mg, 1.2 mmol). The solution was allowed to warm to room temperature after 32 minutes and stirred overnight. The solution was washed two times with brine and dried with magnesium sulfate. The crude product was purified via silica gel chromatography (1% to 10% methanol in DCM) to afford the product in 51% yield as a yellow oil (58 mg).
**$^{1}$H NMR (400MHz, $CDCl_3$)**: δ 6.61-6.53 (m, 1H), 6.35-6.29 (m, 1H), 5.70-5.66 (m, 1H), 3.74-3.72 (m, 1 H), 3.69 (t, *J* = 6.4 Hz, 1H), 3.65-3.61 (m, 2H), 2.66-2.63 (m, 2H), 2.59-2.54 (m, 2H), 2.37 (s, 3H), 1.94 (quint, *J* = 6.2 Hz, 2H).
**$^{13}$C NMR (100MHz, $CDCl_3$)**: δ 166.4, 166.3, 128.0, 127.9, 127.8, 127.6, 59.1, 58.0, 57.1, 56.8, 47.4, 47.1, 46.7, 46.6, 45.3, 44.8, 28.1, 26.9.
**HRMS (+ESI)**: Calculated: 169.1335 ($C_9H_{17}N_2O$). Observed: 169.1333.

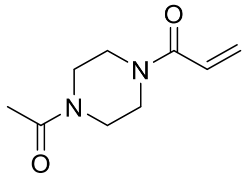

**1-(4-acetylpiperazin-1-yl)prop-2-en-1-one (YP-1-24)**
A solution of 1-acetylpiperazine (128 mg, 1.0 mmol) in DCM (10 mL) was cooled to 0 $^oC$. To the solution was added acryloyl chloride (109 mg, 1.2 mmol) followed by triethylamine (121 mg, 1.2 mmol). The solution was allowed to warm to room temperature after 23 minutes and stirred for two hours. The solution was washed two times with brine and dried with magnesium sulfate. The crude product was purified via silica gel chromatography (0% to 10% methanol in DCM) to afford the product in 18% yield as a yellow oil (40 mg).
**$^1$H NMR (400MHz, $CDCl_3$)**: δ 6.57 (dd, *J* = 10.5, 16.8 Hz, 1H), 6.33 (dd, *J* = 1.8, 16.8 Hz, 1H), 5.75 (dd, *J* = 1.9, 10.5 Hz, 1H), 3.72 (s, 1H), 3.66-3.64 (m, 3H), 3.57 (s, 1H), 3.51-3.49 (m, 2H), 2.13 (s, 3H).
**$^{13}$C NMR (100MHz, $CDCl_3$)**: δ 165.6, 128.7, 127.0, 41.9, 41.4, 21.4.
**HRMS (+ESI)**: Calculated: 183.1128 ($C_9H_{15}N_2O_2$). Observed: 183.1126.

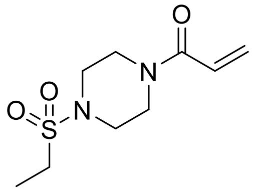

**1-(4-(Ethylsulfonyl)piperazin-1-yl)prop-2-en-1-one (YP-1-25)**
A solution of 1-(ethanesulfonyl)piperazine (178 mg, 1.0 mmol) in DCM (10 mL) was cooled to 0 $^oC$. To the solution was added acryloyl chloride (109 mg, 1.2 mmol) followed by triethylamine (121 mg, 1.2 mmol). The solution was allowed to warm to room temperature after 27 min and stirred for two hours. The solution was washed two times with brine and dried with magnesium sulfate. The crude product was purified via silica gel chromatography (1% to 10% methanol in DCM) to afford the product in 70% yield as a white-yellow solid (163 mg).
**$^1$H NMR (400MHz, $CDCl_3$)**: δ 6.57 (dd, *J* = 10.5, 16.8 Hz, 1H), 6.32 (dd, *J* = 1.9, 16.8 Hz, 1H), 5.76 (dd, *J* = 1.8, 10.5 Hz, 1H), 3.77 (s, 2H), 3.67 (s, 2H), 3.32 (t, *J* = 5.2 Hz, 4H), 2.98 (q, *J* = 7.5 Hz, 2H), 1.37 (t, *J* = 7.4, 3H).
**$^{13}$C NMR (100MHz, $CDCl_3$)**: δ 165.5, 128.8, 127.0, 77.4, 45.9, 45.6, 44.2, 41.9, 7.8.
**HRMS (+ESI)**: Calculated: 233.0954 ($C_9H_{17}N_2O_3S_1$). Observed: 233.0953.

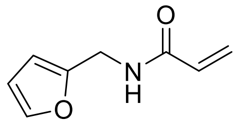

**N-(Furan-2-ylmethyl)acrylamide (YP-1-26)**
A solution of furfurylamine (97 mg, 1.0 mmol) in DCM (10 mL) was cooled to 0 $^oC$. To the solution was added acryloyl chloride (109 mg, 1.2 mmol) followed by triethylamine (121 mg, 1.2 mmol). The solution was allowed to warm to room temperature after 17 min and stirred for two and a half hours. The solution was washed two times with brine and dried with magnesium sulfate. The crude product was purified via silica gel chromatography

(35% to 70% ethyl acetate in hexanes) to afford the product in 86% yield as a white solid (132 mg).
**$^1$H NMR (400MHz, CDCl$_3$)**: δ 7.33 (s, 1H), 6.60 (s, 1H), 6.31-6.22 (m, 3H), 6.15 (dd, *J* = 10.1, 16.9 Hz, 1H), 5.63 (dd, *J* = 1.6, 10.1 Hz, 1H), 4.48 (d, *J* = 5.6 Hz, 2H).
**$^{13}$C NMR (100MHz, CDCl$_3$)**: δ 165.5, 151.2, 142.2, 130.6, 126.8, 110.5, 107.5, 36.5.
**HRMS (+ESI)**: Calculated: 152.0706 ($C_8H_{10}O_2N_1$). Observed: 152.0706.

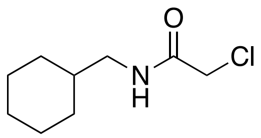


**2-chloro-*N*-(cyclohexylmethyl)acetamide (YP-1-31)**
Following **General Procedure B** starting from cyclohexanemethylamine (113 mg, 1.0 mmol), product was obtained after silica gel chromatography (100% dichloromethane to 3% methanol in dichloromethane) in 60% yield as a white solid (112 mg).
**$^1$H NMR (400MHz, CDCl$_3$)**: δ 6.70 (s, 1H), 4.06 (s, 2H), 3.15 (t, *J* = 6.47 Hz, 2H), 1.77-1.65 (m, 5H), 1.56-1.46 (m, 1H), 1.30-1.10 (m, 3H), 1.00-0.90 (m, 2H).
**$^{13}$C NMR (100MHz, CDCl$_3$)**: δ 165.8, 58.1, 46.0, 42.8, 37.7, 30.7, 26.3, 25.7, 18.2.
**HRMS (+ESI)**: Calculated: 190.0993 ($C_9H_{17}ONCl$). Observed: 190.0992.

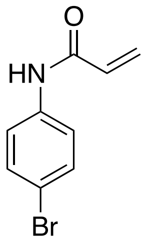


***N*-(4-bromophenyl)acrylamide (YP-1-36)**
Following **General Procedure A** starting from 4-bromoaniline (688 mg, 4.0 mmol), product was obtained after silica gel chromatography (30% to 60% ethyl acetate in hexanes) in 28% yield as a white solid (250 mg).
**$^1$H NMR (400MHz, CD$_3$OD)**: δ 7.90 (s, 1H), 7.60-7.56 (m, 2H), 7.47-7.44 (m, 2H), 6.45-6.33 (m, 2H), 5.78 (dd, *J* = 2.8, 9.1 Hz, 1H).
**$^{13}$C NMR (100MHz, CD$_3$OD)**: δ 164.7, 137.7, 131.4, 130.9, 126.7, 121.5, 116.3, 101.1, 78.1.
**HRMS (+ESI)**: Calculated: 223.9716 ($C_9H_7NOBr$). Observed: 223.9719.

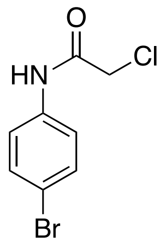


***N*-(4-bromophenyl)-2-chloroacetamide (YP-1-37)**
Following **General Procedure B** starting from 4-bromoaniline (688 mg, 4.0 mmol), product was obtained after silica gel chromatography (30% to 60% ethyl acetate in hexanes) in 49% yield as a white solid (491 mg).

**$^{1}$H NMR (400MHz, $CD_3OD$)**: δ 7.9 (s, 1H), 7.57-7.53 (m, 2H), 7.50-7.47 (m, 2H), 4.17 (s, 2H).
**$^{13}$C NMR (100MHz, $CD_3OD$)**: δ 166.0, 137.2, 131.5, 121.6, 116.7, 99.3, 78.1, 42.6.
**HRMS (+ESI)**: Calculated: 245.9327 ($C_8H_6NOBrCl$). Observed: 245.9329.

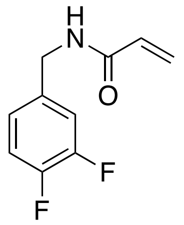

***N*-(3,4-difluorobenzyl)acrylamide (YP-1-38)**
Following **General Procedure A** starting from 3,4-difluorobenzylamine (286 mg, 2.0 mmol), product was obtained after silica gel chromatography (40% to 80% ethyl acetate in hexanes) in 61% yield as a white solid (239 mg).
**$^{1}$H NMR (400MHz, $CDCl_3$)**: δ 7.56 (t, *J* = 6.2, 1H), 7.07-7.00 (m, 2H), 6.95-6.91 (m, 1H), 6.21-6.20 (m, 2H), 5.62-5.59 (m, 1H), 4.35 (d, *J* = 6.1, 2H).
**$^{13}$C NMR (100MHz, $CDCl_3$)**: δ 166.1, 151.4 (d), 150.7 (d), 148.9 (d), 148.2 (d), 135.5-135.4 (m), 130.5, 126.8, 123.5-123.4 (m), 117.2 (d), 116.3 (d), 42.4.
**HRMS (+ESI)**: Calculated: 196.0579 ($C_{10}H_8NOF_2$). Observed: 196.0582.

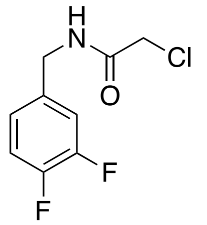

**2-chloro-*N*-(3,4-difluorobenzyl)acetamide (YP-1-39)**
Following **General Procedure B** starting from 3,4-difluorobenzylamine (286 mg, 2.0 mmol), product was obtained after silica gel chromatography (40% to 50% ethyl acetate in hexanes) in 82% yield as a white solid (359 mg).
**$^{1}$H NMR (400MHz, $CDCl_3$)**: δ 7.23 (s, 1H), 7.15-7.08 (m, 2H), 7.03-7.6.99 (m, 1H), 4.42 (d, *J* = 6.1 Hz, 2H), 4.08 (s, 2H).
**$^{13}$C NMR (100MHz, $CDCl_3$)**: δ 166.3, 151.5 (d), 151.0 (d), 149.1 (d), 148.5 (d), 134.7-134.6 (m), 123.7-123.6 (m), 117.5 (d), 116.6 (d), 42.6 (d).
**HRMS (+ESI)**: Calculated: 218.0190 ($C_9H_7NOClF_2$). Observed: 218.0192

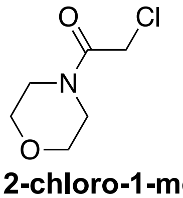

**2-chloro-1-morpholinoethan-1-one (YP-1-40)**
Following **General Procedure B** starting from morpholine (174 mg, 2.0 mmol), product was obtained after silica gel chromatography (85% ethyl acetate in hexanes) in 61% yield as a white solid (200 mg).
**$^{1}$H NMR (400MHz, $CDCl_3$)**: δ 4.01 (s, 2H), 3.65-3.59 (m, 4H), 3.55-3.52 (m, 2H), 3.45 (t, *J* = 4.8 Hz, 2H).
**$^{13}$C NMR (100MHz, $CDCl_3$)**: δ 165.1, 66.5 (d), 46.6, 42.4, 40.7.

**HRMS (+ESI)**: Calculated: 186.0292 ($C_6H_{10}O_2NClNa$). Observed: 186.0292.

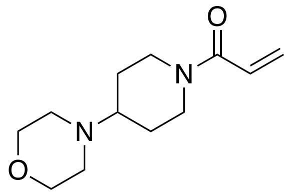

**1-(4-morpholinopiperidin-1-yl)prop-2-en-1-one (YP-1-42)**
Following **General Procedure A** starting from 4-morpholinopiperidine (336 mg, 2.0 mmol), product was obtained after silica gel chromatography (1% methanol and 80% ethyl acetate in hexanes) in 58% yield as a colorless oil (259 mg).
**$^1$H NMR (400MHz, CDCl$_3$)**: δ 6.42 (dd, *J* = 10.6, 16.8 Hz, 1H), 6.06 (dd, *J* = 2.0, 16.8 Hz, 1H), 5.49 (dd, *J* = 2.0, 10.6 Hz, 1H), 4.45 (d, *J* = 12.8 Hz, 1H), 3.86 (d, *J* = 12.8 Hz, 1H), 3.52 (t, *J* = 4.7 Hz, 4H), 2.90 (t, *J* = 12.8 Hz, 1H), 2.55-2.48 (m, 1H), 2.37-2.35 (m, 4H), 2.26 (tt, *J* = 3.7, 11.0 Hz, 1H), 1.72 (d, *J* = 12.8 Hz, 2H), 1.30-1.20 (m, 2H).
**$^{13}$C NMR (100MHz, CDCl$_3$)**: δ 165.0, 127.7, 127.3, 67.1, 61.6, 49.6, 44.9, 41.1, 28.9, 27.8.
**HRMS (+ESI)**: Calculated: 225.1598 ($C_{12}H_{21}N_2O_2$). Observed: 225.1595.

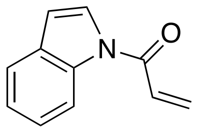

**1-(1*H*-indol-1-yl)prop-2-en-1-one (YP-1-44)**
A solution of indole (117 mg, 1.0 mmol) in 2-methyltetrahydrofuran (10 mL) was cooled to 0°C. To the solution was added sodium hydride (60 mg, 2.5 mmol). The resultant intermediate was subjected to **General Procedure A** and product was obtained after alumina gel chromatography (10% to 40% ethyl acetate in hexanes) in 8% yield as a white solid (14 mg).
**$^1$H NMR (400MHz, CD$_3$OD)**: δ 8.48-8.46 (m, 1H), 7.82 (d, *J* = 3.9 Hz, 1H), 7.61-7.59 (m, 1H), 7.36-7.32 (m, 1H), 7.31-7.21 (m, 2H), 6.73 (dd, *J* = 0.8, 3.8 Hz, 1H), 6.64 (dd, *J* = 1.7, 16.7 Hz, 1H), 6.09 (dd, *J* = 1.7, 10.5 Hz, 1H).
**$^{13}$C NMR (100MHz, CD$_3$OD)**: δ 164.3, 135.7, 131.0, 130.9, 128.0, 125.0, 124.4, 123.6, 120.5, 116.2, 108.9.
**HRMS (+ESI)**: Calculated: 172.0757 ($C_{11}H_{10}NO$). Observed: 172.0756.

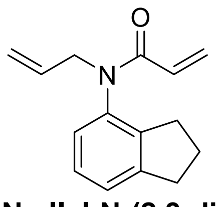

**N-allyl-N-(2,3-dihydro-1H-inden-4-yl)acrylamide (IGA-1-12)**
A solution of sodium hydride (96 mg, 4.0 mmol) in tetrahydrofuran (8 mL) was put under nitrogen atmosphere. To the solution was added N-(2,3-dihydro-1H-inden-4-yl)acrylamide (187 mg, 1.0 mmol) in tetrahydrofuran (2 mL). The solution was cooled to 0 °C and stirred. 3-bromoprop-1-ene (484 mg, 4.0 mmol) was added after 30 minutes, after which the solution was allowed to warm to room temperature and was stirred overnight. The solution was quenched with water and extracted with ethyl acetate. The

crude product was purified via silica gel chromatography (20% ethyl acetate in hexanes) to afford the product in 67% yield as a yellow crystalline solid (151 mg).
**$^{1}$H NMR (400MHz, $CDCl_3$)**: δ 7.06-7.18 (m, 2H), 6.80-6.88 (m, 1H), 6.26-6.37 (dd, *J* = 16.8, 2.0 Hz, 1H), 5.76-5.96 (m, 2H), 5.38-5.48 (dd, *J* = 10.3, 2.1 Hz, 1H), 4.98-5.08 (m, 2H), 4.40-4.52 (ddt, *J* = 14.5, 6.3, 1.3 Hz, 1H), 4.00-4.11 (ddt, *J* = 14.5, 6.8, 1.2 Hz, 1H), 2.82-2.98 (m, 2H), 2.59-2.79 (m, 2H), 1.92-2.07 (m, 2H).
**$^{13}$C NMR (100MHz, $CDCl_3$)**: δ 165.1, 146.5, 142.4, 137.9, 133.0, 128.4, 127.8, 127.48, 126.1, 124.3, 118.1, 51.6, 33.3, 30.9, 25.0.
**HRMS (+ESI)**: Calculated: 228.13 ($C_{15}H_{17}NO$). Observed: 228.1381.

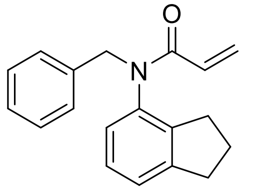

**N-benzyl-N-(2,3-dihydro-1H-inden-4-yl)acrylamide (IGA-1-14)**
A solution of sodium hydride (96 mg, 4.0 mmol) in tetrahydrofuran (8 mL) was put under nitrogen atmosphere. To the solution was added N-(2,3-dihydro-1H-inden-4-yl)acrylamide (187 mg, 1.0 mmol) in tetrahydrofuran (2 mL). The solution was cooled to 0 °C and stirred. Benzyl bromide (476 mg, 4.0 mmol) was added after 30 minutes, after which the solution was allowed to warm to room temperature and was stirred overnight. The solution was quenched with water and extracted with ethyl acetate. The crude product was purified via silica gel chromatography (20% ethyl acetate in hexanes) to afford the product in 63% yield as an orange oil (173 mg).
**$^{1}$H NMR (400MHz, $CDCl_3$)**: δ 7.10-7.35 (m, 7H), 6.74-6.85 (dd, *J* = 7.8, 1.1 Hz, 1H), 6.40-6.55 (dd, *J* = 16.8, 2.1 Hz, 1H), 5.93-6.08 (dd, *J* = 16.8, 10.3 Hz, 1H), 5.49-5.62 (dd, *J* = 10.3, 2.1 Hz, 1H), 4.78-5.10 (m, 2H), 2.85-3.02 (m, 2H), 2.52-2.67 (m, 1H), 2.22-2.37 (m, 1H), 1.83-2.01 (m, 2H).
**$^{13}$C NMR (100MHz, $CDCl_3$)**: δ 146.4, 142.9, 137.7, 137.3, 129.3, 128.4, 128.3, 128.0, 127.5, 127.5, 126.0, 124.3, 52.3, 33.2, 30.6, 25.1.
**HRMS (+ESI)**: Calculated: 278.15 ($C_{19}H_{19}NO$). Observed: 278.1538.

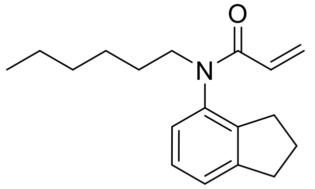

**N-allyl-N-(2,3-dihydro-1H-inden-4-yl)acrylamide (IGA-1-15)**
A solution of sodium hydride (96 mg, 4.0 mmol) in tetrahydrofuran (8 mL) was put under nitrogen atmosphere. To the solution was added N-(2,3-dihydro-1H-inden-4-yl)acrylamide (187 mg, 1.0 mmol) in tetrahydrofuran (2 mL). The solution was cooled to 0 °C and stirred. 1-bromohexane (660 mg, 4.0 mmol) was added after 30 minutes, after which the solution was allowed to warm to room temperature and was stirred overnight. The solution was quenched with water and extracted with ethyl acetate. The crude product was purified via silica gel chromatography (20% ethyl acetate in hexanes) to afford the product in 34% yield as a yellow oil (92 mg).
**$^{1}$H NMR (400MHz, $CDCl_3$)**: δ 7.11-7.25 (m, 2H), 6.86-6.96 (dd, *J* = 7.5, 1.2 Hz, 1H), 6.30-6.40 (dd, *J* = 16.8, 2.1 Hz, 1H), 5.86-6.00 (m, 1H), 5.41-5.51 (dd, *J* = 10.3, 2.1 Hz, 1H), 3.82-3.96 (m, 1H), 3.42-3.56 (m, 1H), 2.90-3.04 (m, 2H), 2.65-2.85 (m, 2H), 1.98-2.16 (m, 2H), 1.47-1.63 (m, 2H), 1.20-1.36 (m, 6H), 0.80-0.90 (m, 3H).

**$^{13}$C NMR (100MHz, $CDCl_3$)**: δ 165.16, 146.54, 142.38, 138.21, 128.59, 127.51, 127.35, 126.09, 124.13, 48.67, 33.26, 31.62, 30.85, 27.85, 26.72, 25.01, 22.59, 14.05.
**HRMS (+ESI)**: Calculated: 272.19 ($C_{18}H_{25}NO$). Observed: 272.2007.

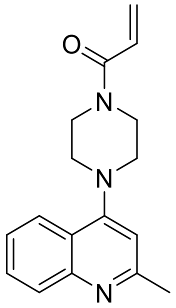


**1-(4-(2-methylquinolin-4-yl)piperazin-1-yl)prop-2-en-1-one (IGA-1-26)**
A solution of 2-methyl-4-(piperazin-1-yl)quinolone (455 mg, 2.0 mmol) in DCM (20 mL) was cooled to 0 °C. To the solution was added acryloyl chloride (217 mg, 2.4 mmol) followed by triethylamine (243 mg, 2.4 mmol). The solution was allowed to warm to room temperature and stirred overnight. The solution was washed with brine and the crude product was purified via basic alumina chromatography (100% ethyl acetate) to afford the product in 26% yield as a yellow oil (145 mg).
**$^{1}$H NMR (400MHz, $CDCl_3$)**: δ 7.90-8.05 (m, 2H), 7.58-7.70 (ddd, $J$ = 8.4, 6.8, 1.5 Hz, 1H), 7.40-7.50 (ddd, $J$ = 8.2, 6.8, 1.3 Hz, 1H), 6.68-6.76 (s, 1H), 6.56-6.67 (dd, $J$ = 16.8, 10.5 Hz, 1H), 6.30-6.40 (dd, $J$ = 16.8, 2.0 Hz, 1H), 5.70-5.80 (dd, $J$ = 10.5, 2.0 Hz, 1H), 3.70-4.06 (d, $J$ = 54.7 Hz, 4H), 3.10-3.30 (t, $J$ = 5.0 Hz, 4H), 2.62-2.72 (s, 3H).
**$^{13}$C NMR (100MHz, $CDCl_3$)**: δ 165.5, 159.4, 156.2, 149.2, 129.26, 129.24, 128.3, 127.3, 124.9, 123.0, 121.6, 109.8, 52.3, 51.9, 45.8, 42.0, 25.6.
**HRMS (+ESI)**: Calculated: 282.17 ($C_{17}H_{19}N_3O$). Observed: 282.1597.

**Supporting Table Legends**

**Table S1. IsoTOP-ABPP analysis of parthenolide targets.** List of probe-modified peptides, protein designations, and their light to heavy ratios for those peptides identified in two out of four biological replicates. 231MFP breast cancer cell proteomes were treated with DMSO vehicle or parthenolide (50 μM) for 30 min followed by labeling with IA-alkyne (100 μM) for 1 h. Probe-labeled proteins were then subjected to CuAAC-mediated appendage of a biotin-azide tag bearing a TEV protease recognition site and an isotopically light (for control) or heavy (for parthenolide-treated) valine. Control and treated proteomes were combined in a 1:1 ratio and probe-modified proteins were avidin-enriched and tryptically digested. Probe-modified tryptic peptides were subsequently avidin-enriched and eluted using TEV protease. Probe-modified peptides were analyzed by LC-MS/MS and light and heavy peptides were quantified. Shown are average ratios for each peptide from n=4. **Table S1** is related to **Figure 2.**

**Table S2**. **Structures of cysteine-reactive covalent ligands screened against FAK1. Table S2** is related to **Figure S2-S4**.

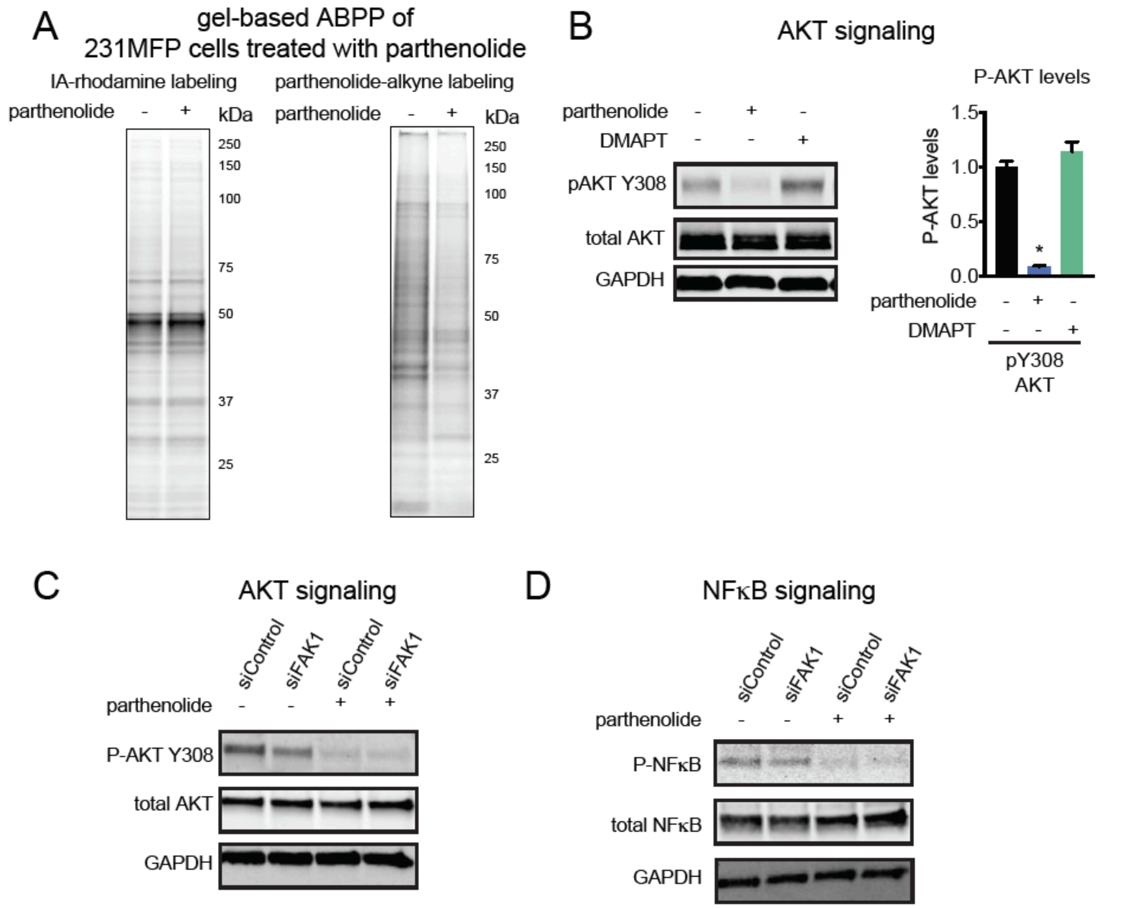


**Figure S1. Effects of parthenolide. (A)** Gel-based ABPP analysis of parthenolide in 231MFP proteomes *in situ*. In the left panel, 231MFP cells were pre-incubated with parthenolide (50 μM) for 30 min prior to labeling with IA-rhodamine (1 μM) for 1 h, after which cells were harvested and subjected to SDS/PAGE and in-gel fluorescence. In the right panel, 231MFP cells were pre-incubated with parthenolide (50 μM) for 30 min prior to parthenolide-alkyne labeling (50 μM) for 1 h, after which rhodamine-azide was appended by CuAAC, and proteome was separated by SDS/PAGE and visualized by in-gel fluorescence. **(B)** AKT signaling in 231MFP cells treated with vehicle DMSO, parthenolide (50 μM), or DMAPT (50 μM) for 2 h, assessed by Western blotting. **(C, D)** AKT **(C)** and NFκB **(D)** signaling in 231MFP siControl and siFAK1 cells treated with vehicle DMSO or parthenolide (50 μM) for 2 h. Gels shown in **(A-D)** are representative gels of n=3 biological replicates/group. Bar graph shown in **(B)** is average ± sem, n=3/group.

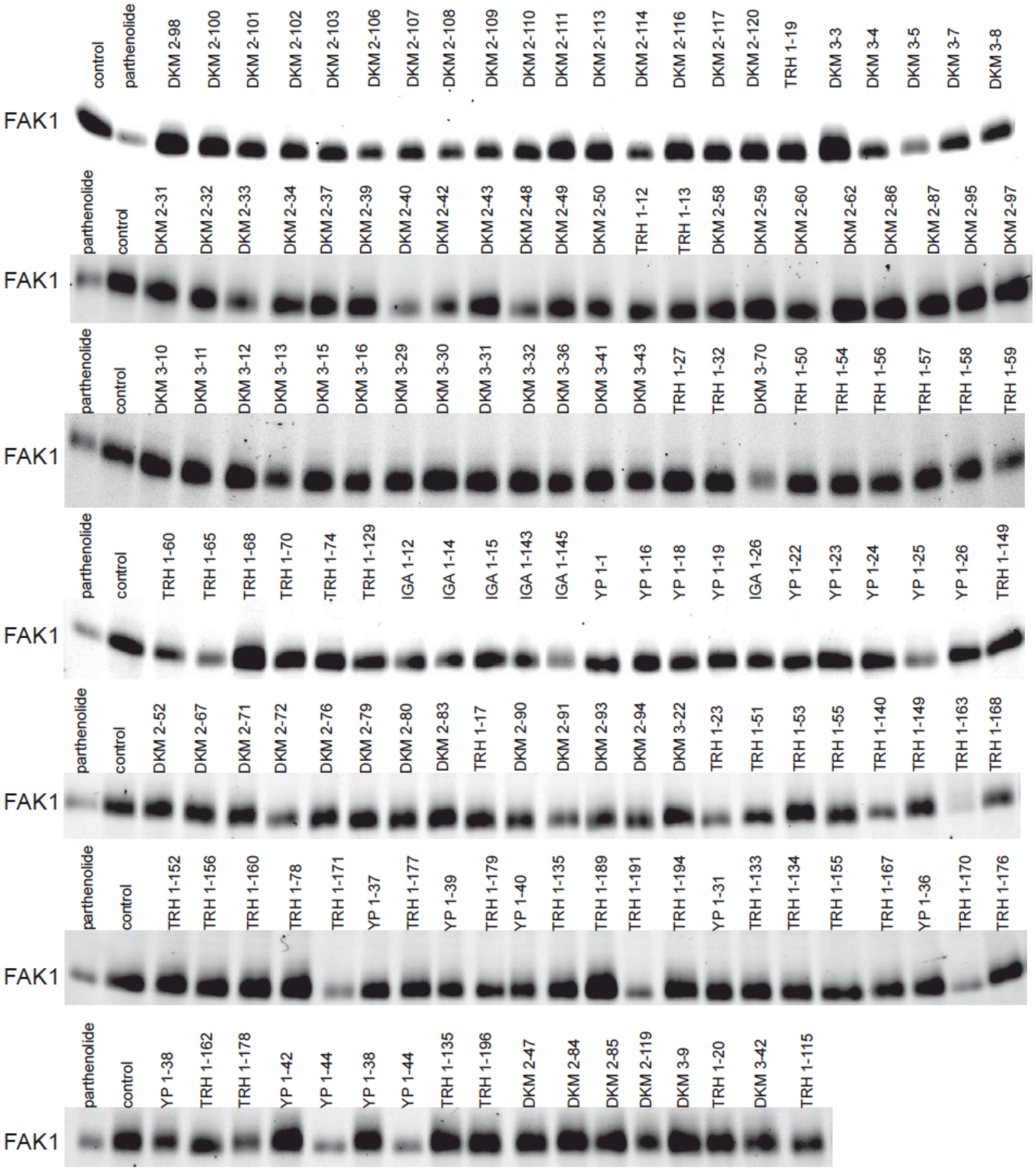


**Figure S2. Covalent ligand screen against FAK1**. Gel-based ABPP screening of cysteine-reactive fragment library (50 $\mu$M, 30 min pre-incubation) against IA-rhodamine labeling (1 $\mu$M, 1  labeling). Probe-labeled proteins were analyzed by in-gel fluorescence. **Figure S2** is related to **Figure 4**.

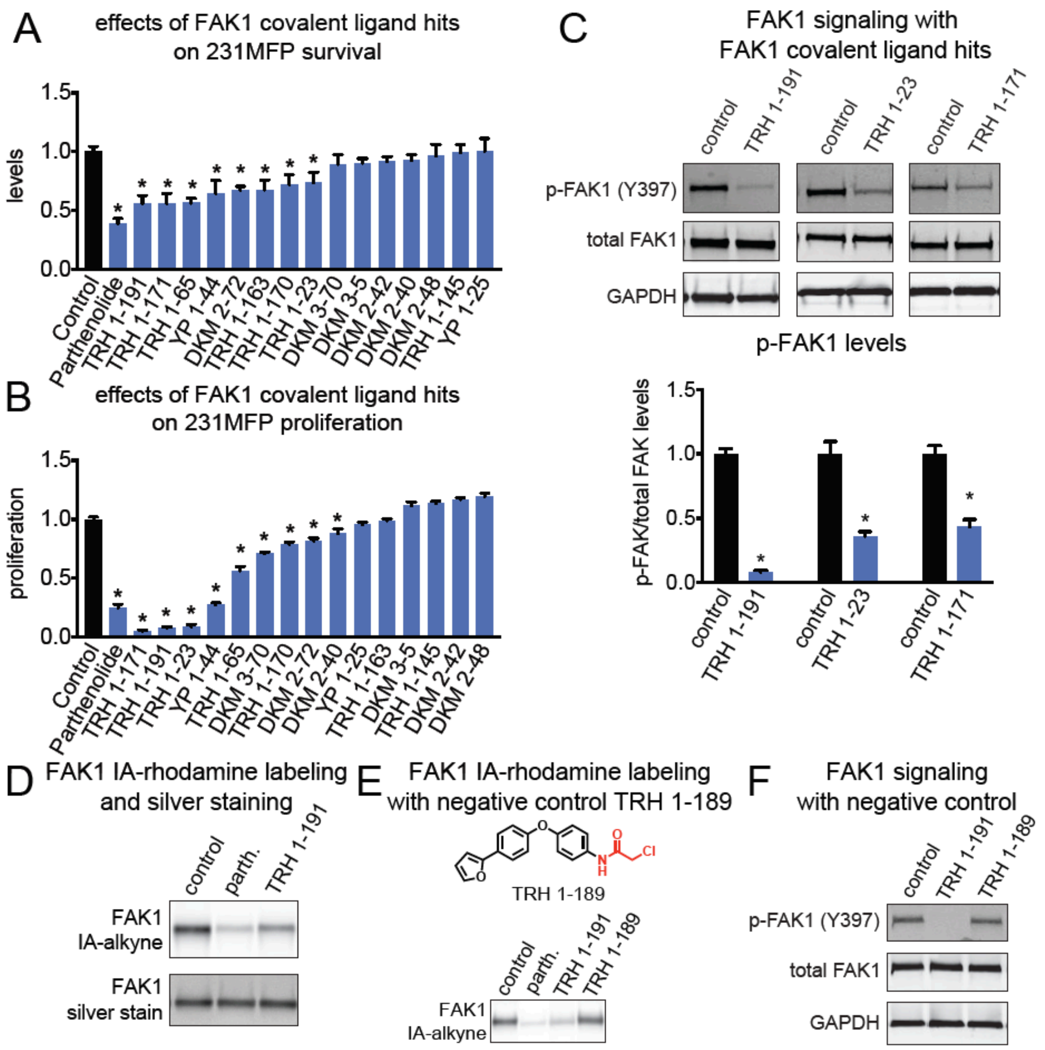


**Figure S3. Effects of FAK1 covalent ligands on breast cancer cells. (A, B)** 231MFP breast cancer cell survival and proliferation (48 h) from cells treated with DMSO vehicle, parthenolide, or FAK1 covalent ligand hits (50 μM), assessed by Hoechst stain. **(C)** FAK1 signaling in breast cancer cells treated with DMSO vehicle or FAK1 covalent ligand hits (50 μM). Gel images are representative images from n=3. **(D)** Gel-based ABPP and silver staining analysis of parthenolide and TRH 1-191 *in vitro* treatment with human FAK1 pure protein. FAK1 protein was pre-incubated with parthenolide or TRH 1-191 (50 μM, 30 min) prior to IA-rhodamine labeling (11 μM, 1 h) and in-gel fluorescence analysis. Gels were also silver stained to confirm equal FAK1 protein loading. **(E)** Gel-based ABPP analysis of TRH 1-191 analog TRH 1-189 that does not inhibit FAK1 IA-alkyne labeling. **(F)** FAK1 signaling assessed by Western blotting. 231MFP cells were treated with DMSO vehicle or TRH 1-191 or TRH 1-189 (50 μM) for 2 h. **(G)** 231MFP breast cancer cell survival and proliferation (48 h) from cells treated with DMSO vehicle, TRH 1-191, or TRH 1-189 (50 μM), assessed by Hoechst stain. Data shown are average ± sem, n=3-6/group. Significance is expressed as *p<0.05 compared to vehicle-treated controls. **Figure S3** is related to **Figure S2**.

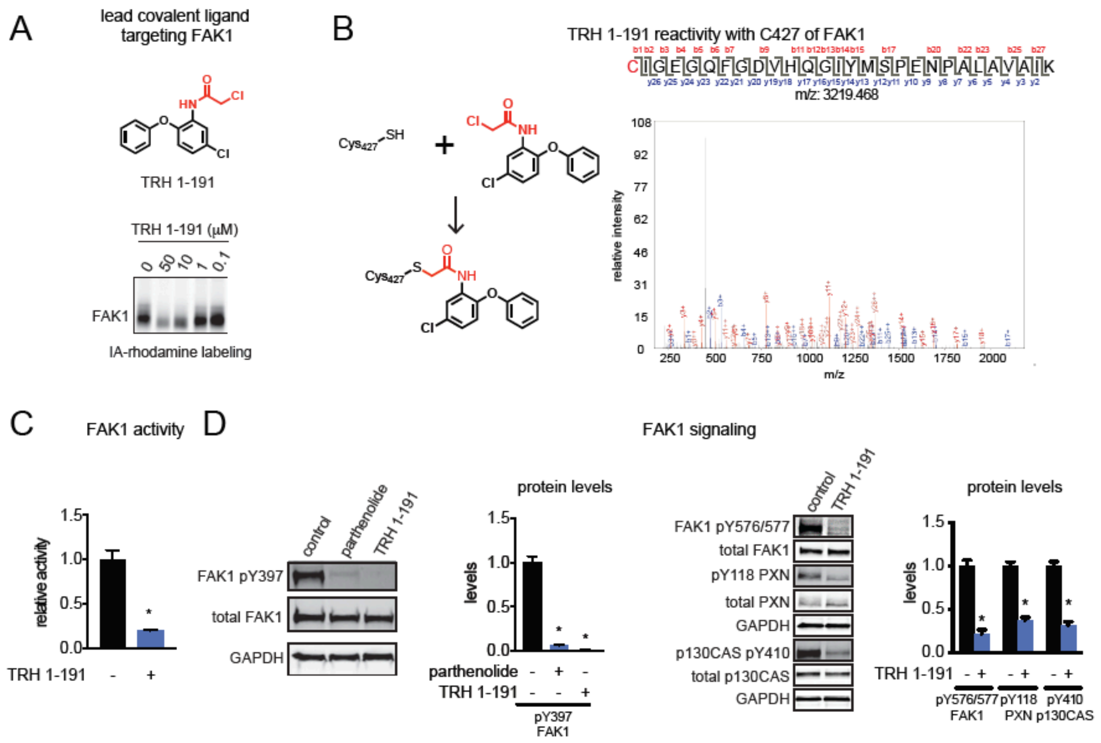


**Figure S4. Covalent ligand TRH 1-191 reacts with C427 of FAK1 and inhibits FAK1 signaling and breast cancer pathogenicity. (A)** Structure of TRH 1-191 showing the cysteine-reactive chloroacetamide warhead in red. Shown below is a gel-based ABPP analysis of TRH 1-191 competition against IA-rhodamine labeling of pure human FAK1 protein. **(B)** Presumed reaction of TRH 1-191 with C427 FAK1 and mass spectrometry data showing the TRH 1-191 adduct on C427 of FAK1. Pure human FAK1 kinase domain was treated with TRH 1-191 (100 µM) and the protein was subsequently digested with trypsin for LC-MS/MS analysis. **(C)** FAK1 activity of pure human FAK1 kinase domain assessed by peptide substrate phosphorylation and read-out by ADP-Glo kinase assay. FAK1 protein was treated with DMSO vehicle or TRH 1-191 (100 µM) for 30 min prior to addition of substrates. **(D)** FAK1 signaling assessed by Western blotting in 231MFP cells treated with vehicle DMSO or TRH 1-191 (50 µM) for 2 h. Gels shown in **(A, D)** are representative of n=3. Data shown in **(C-D)** are average ± sem, n=3 biological replicates/group. Significance is expressed as *p<0.05 compared to vehicle-treated controls. **Figure S4** is related to **Table S2** and **Figure S2** and **Figure S3**.
